# Model-guided design of microRNA-based gene circuits supports precise dosage of transgenic cargoes into diverse primary cells

**DOI:** 10.1101/2024.06.25.600629

**Authors:** Kasey S. Love, Christopher P. Johnstone, Emma L. Peterman, Stephanie Gaglione, Kate E. Galloway

## Abstract

To realize the potential of engineered cells in therapeutic applications, transgenes must be expressed within the window of therapeutic efficacy. Differences in copy number and other sources of extrinsic noise generate variance in transgene expression and limit the performance of synthetic gene circuits. In a therapeutic context, supraphysiological expression of transgenes can compromise engineered phenotypes and lead to toxicity. To ensure a narrow range of transgene expression, we design and characterize **Co**mpact **m**icroRNA-**M**ediated **A**ttenuator of **N**oise and **D**osage (**ComMAND**), a single-transcript, microRNA-based incoherent feedforward loop. We experimentally tune the ComMAND output profile, and we model the system to explore additional tuning strategies. By comparing ComMAND to two-gene implementations, we highlight the precise control afforded by the single-transcript architecture, particularly at relatively low copy numbers. We show that ComMAND tightly regulates transgene expression from lentiviruses and precisely controls expression in primary human T cells, primary rat neurons, primary mouse embryonic fibroblasts, and human induced pluripotent stem cells. Finally, ComMAND effectively sets levels of the clinically relevant transgenes FMRP1 and FXN within a narrow window. Together, ComMAND is a compact tool well-suited to precisely specify expression of therapeutic cargoes.

## Introduction

With massive advances in the development of delivery vectors, the tunable control of therapeutic cargoes remains the missing element in providing safe, effective *in vivo* and *ex vivo* gene therapies [1–9]. A large number of diseases result from mutations affecting a single gene [10]. Many of these monogenic disorders render the cell vulnerable by the insufficient production of an essential protein product [11–15]. For these haploinsufficiencies, delivery of functional copies of affected genes may restore essential cellular processes and rescue normal phenotypes. However, while improved vectors can efficiently deliver synthetic cargoes, excessive expression of transgenes can induce unforeseen neurological and cardiac disorders. For instance, in mouse models of neurological disorders that result from haploinsufficiency, gene replacement therapy generated mixed outcomes. When target proteins were overexpressed at ten to twenty times their endogenous level, some mice showed improved function, but others suffered adverse events including cardiotoxicity and behavioral abnormalities [16–18]. For safety and efficacy, delivery of therapeutic cargoes must be tailored to ensure that transgenes are expressed within a “goldilocks” window (Figure 1A). Thus, there is a need for tools that can precisely set levels of transgene expression.

**Figure 1.**
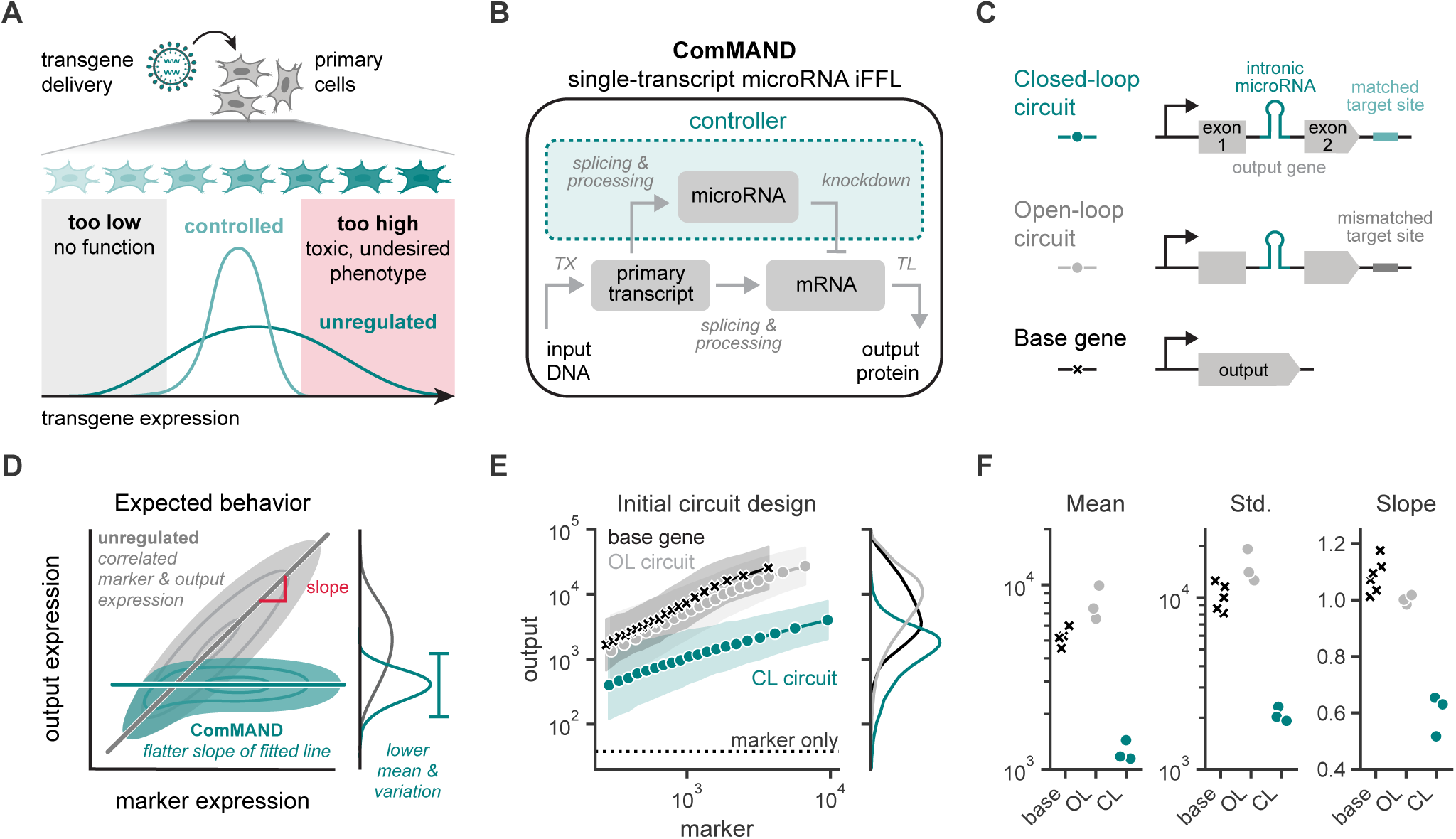
ComMAND, a single-transcript, microRNA-based incoherent feedforward loop (iFFL), reduces output mean and variability compared to unregulated genes. **A.** Delivery of transgenes to primary cells results in high expression variance. Unregulated transgene expression levels may be too low, where the transgene does not function, or too high, which can lead to toxicity or other undesired phenotypes. Controlled transgene expression has the potential to narrow this expression distribution to within the functional range. **B.** Block diagram of ComMAND, a single-transcript, microRNA-based iFFL. DNA, the input, is transcribed (TX) into a primary transcript, which is then spliced and processed into mature mRNA and microRNA. The microRNA can knock down the mRNA; microRNA production and function constitute the controller (teal). Alternatively, the mRNA can be translated (TL) into protein, the output of the circuit. **C.** DNA construct diagrams of ComMAND. The closed-loop (CL) circuit consists of an intronic microRNA between two exons of the output gene and a 22-bp complementary microRNA target site in the 3’UTR. In the open-loop (OL) circuit, the microRNA target site is orthogonal to the microRNA sequence. The base gene includes only the output coding sequence, lacking both the microRNA and the target site. **D.** Expected behavior of ComMAND across a population of cells when co-delivered with a constitutively expressed marker gene. For an unregulated gene (base gene or OL circuit), the marker and output expression should be well-correlated, giving rise to a wide output expression distribution with a positive slope near one (gray). In contrast, the closed-loop circuit is expected to display relatively constant output expression levels across a range of marker expression, leading to a lower output expression mean, a narrower output distribution, and a flatter slope (teal). **E.** Left: Output expression as a function of marker expression for constructs in Figure 1C co-transfected with a marker gene in HEK293T cells. Flow cytometry measurements for one representative biological replicate are binned by marker expression into 20 equal-quantile groups per condition. Points represent geometric means of output expression for cells in each bin, and shaded regions represent this value multiplied or divided by the geometric standard deviation of the bin. Bins are plotted at their median marker value. Dashed line represents the output geometric mean of cells transfected only with the marker gene. Right: Histograms of output expression for cells in each condition. The output gene, mRuby2, was expressed via the EF1α promoter, and circuits included miR-FF5. **F.** Summary statistics of output expression for populations in 1E. The plotted mean values use the geometric mean. Std. refers to the standard deviation, and the slope represents the slope of the line fitted to the binned, log-transformed marker-output points in 1E. Points represent individual biological replicates. All units are arbitrary units (AU) from a flow cytometer.

Gene dosage, including gene copy number, represents a major source of variance in gene expression [19]. While efficient at delivering genetic cargoes, random integration methods offer limited control of copy number. Adjusting vector dosage simultaneously—and unavoidably—affects both delivery efficiency and copy number, making it difficult to maintain high coverage of cells while also ensuring most cells receive only one or a few copies of the transgene. Uneven biodistribution may further compound the challenge of tailoring expression by adjusting dosage [20]. Furthermore, for integrating vectors, differences in the site of integration introduce distinct genomic contexts for transgenes, which can generate cell-to-cell variability in transcription and thus expression [21]. Combined with wide variation in promoter activity across cell types [22, 23], local effects on transcriptional activity present challenges to robust, precise transgene expression.

Gene circuits offer a solution for implementing tight control of transgene expression. Gene circuits such as the incoherent feedforward loop (iFFL) can reduce expression variation caused by DNA copy number [24–30]. To do so, the iFFL senses and compensates for changes in the input by both positively (directly) and negatively (indirectly via an additional species) regulating the output. Mathematical analysis of the iFFL demonstrates that this topology—in the absence of resource limitations—enables perfect adaptation, in which the system response returns to its original level after a disturbance [31–36]. Previous work has implemented iFFLs in mammalian cells via transcriptional [25, 37] and post-transcriptional [1, 25–28, 30, 33] control and demonstrated experimentally that this circuit architecture enables robustness to resource competition and sources of extrinsic noise such as plasmid copy number. A similar circuit has also been used to mitigate toxic transgene expression during AAV production [38]. Constructing an iFFL using microRNA elements is particularly advantageous because these components are small and may require fewer cellular resources to function. However, until recently [1, 30], it remained unclear if a microRNA-based implementation of this circuit could reduce variability introduced by random integration methods or function effectively in primary cells. Furthermore, there is a need to articulate design rules to build circuits that function across diverse delivery methods and to identify optimal performance regimes.

To this end, we constructed a genetic control system called **Co**mpact **m**icroRNA-**M**ediated **A**ttenuator of **N**oise and **D**osage (**ComMAND**). ComMAND comprises a microRNA-based iFFL encoded on a single transcript (Figure 1B). Instead of relying on protein components, ComMAND requires only the addition of an intronic microRNA and a corresponding target site, allowing these circuits to remain extremely compact with minimal resource burden and low immunogenicity. The single-transcript architecture requires fewer genetic elements than multi-transcript or multi-vector designs, making it advantageous for use in therapeutic applications where delivery poses a key challenge. Furthermore, encoding the microRNA in an intron of the output gene closely links microRNA and output mRNA production, since transcription and proper splicing are required to produce molecules of each of these species. The single-transcript design thus more tightly couples the direct and indirect effects of the iFFL on the output, affording more precise control.

Additionally, this intronic design provides an inherently fail-safe mechanism of control. Unlike non-intronic designs, if splicing of the microRNA fails, the regulated transgene cannot be correctly translated from that transcript. Finally, because ComMAND functions downstream of transcription, it is compatible with a wide range of expression methods, including cell type-specific or small molecule-inducible promoters, enabling additional layers of regulation. Here, we characterize ComMAND and demonstrate its generalizability across cell types, delivery methods, and regulated genes, paving the way for therapeutic applications.

## Results

### A single-transcript, microRNA-based incoherent feedforward loop (iFFL) reduces the expression mean and variability of an output protein

To take advantage of the compact size of microRNA components, we constructed ComMAND, a single-transcript version of an incoherent feedforward loop (iFFL) (Figure 1B). To prevent crosstalk with endogenous transcripts, we selected a panel of previously developed synthetic microRNAs and cognate target sequences derived from firefly luciferase [39]. These microRNAs are expressed within an intron containing the human miR-30a scaffold and have targeting sequences orthogonal to the human genome. We first verified that this panel of microRNAs efficiently knocks down their targets in transfections of HEK293T cells (Figure S1A). Of the four reported sequences, we found that three sequences (FF4, FF5, and FF6) are orthogonal and effective at target knockdown (Figures S1B and S1C). Additionally, presence of the microRNA or target site alone did not alter gene expression (Figures S1D and S1E). To construct the iFFL, we inserted an intron bearing the FF5 microRNA (miR-FF5) at an AGGT sequence within the output gene, mRuby2, to generate favorable splice donor and acceptor sites. The intronic design of the iFFL adds only *∼*450 bp to the total transcript size, ensuring ComMAND remains sufficiently compact for therapeutic applications with strict cargo limits. For the closed-loop (CL) circuit, we included a perfectly complementary 22-bp FF5 target sequence in the 3’UTR of the transcript (Figure 1C). For comparison, we constructed an open-loop (OL) version of the circuit that replaces the complementary target site with one of the other orthogonal sequences from our panel. We also included a “base gene”: a second unregulated control that lacks both the microRNA-containing intron and the microRNA target site (i.e., the output gene alone). Together, these constructs enable us to parse the effects of the circuit on output expression.

To examine the ability of ComMAND to control variation introduced by differences in copy number and other extrinsic factors, we co-delivered the circuit with a separate fluorescent marker gene. Expression of the fluorescent marker varies across a population of cells as a function of DNA dosage and cellular physiology. For unregulated genes, the output and marker expression should be strongly correlated (Figure 1D, gray). We quantify correlation by the slope of the marker-output line in logarithmic space, which indicates how much the output expression varies as marker expression changes. A slope below one indicates sublinear scaling. For unregulated genes, we expect marker and output expression to co-vary, resulting in a slope around one. In contrast, we expect that the CL circuit will have relatively lower output expression variation across a range of marker levels (Figure 1D, teal). As ComMAND approaches optimal performance, the slope will approach zero. The CL circuit should also have a lower mean output level than an unregulated gene due to knockdown of the output mRNA. Indeed, when we co-transfected the base gene, OL circuit, or CL circuit with a marker gene in HEK293T cells, we observed a reduction in output expression level, output variation, and slope of the marker-output line for the CL circuit compared to the unregulated genes (Figures 1E and 1F). ComMAND significantly reduces the variability of the output gene and achieves sublinear scaling but does not provide perfect adaptation to extrinsic noise. To better understand the system and improve performance, we next examined how to tune circuit output.

### Selection of circuit components enables tuning of output expression

In order to define design principles for implementing ComMAND, we characterized output expression profiles of constructs with different sets of genetic parts. We sought to understand how part identity contributes to circuit performance and to determine methods to set output expression levels. First, we investigated how properties of the microRNA affect output expression. Increasing the strength of the repression arm of the iFFL should reduce output variability [28]. As FF4 showed the greatest target knockdown (Figure S1B), we hypothesized that replacing the FF5 sequences with FF4 sequences in ComMAND would improve regulation of the output gene. Indeed, introduction of FF4 reduces output mean and standard deviation for the CL circuit in transfection of HEK293T cells (Figures S1H, 2A and 2C). To further increase knockdown, we next modified the microRNA scaffold. Previously, rational sequence modifications to enhance pri-microRNA processing led to the development of the Mv3 (miRE) scaffold, which improves AND gate function in transfection of HEK293T cells [40]. Therefore, we replaced the miR-30a–based scaffold with the miRE scaffold in ComMAND. The miRE scaffold improves the performance of the FF5 CL circuit, approaching the performance of the FF4 CL circuit (Figures S1H, 2B and 2C). However, both miRE-FF4 and miRE-FF5 in ComMAND perform similarly to the miR-FF4 design. Because miRE-FF4 putatively has the strongest target knockdown, we moved forward with miRE-FF4 in ComMAND.

Since altering the microRNA sequence had only a small effect on output expression profiles, we considered other methods to tune the circuit. As ComMAND acts at the post-transcriptional level (Figure 1B), it is compatible with diverse promoters. Tuning promoter strength can alter the setpoint of microRNA-based iFFLs [30], but it remains unclear how this property affects single-transcript architectures. To explore how promoter selection affects ComMAND output expression, we transfected circuits driven by a panel of strong and weak promoters (Figure 2D). Output levels are higher with the strong promoters compared to with the weaker promoters as expected. In all cases, the CL circuit reduces expression level and variability compared to the base gene and OL circuit with the same promoter (Figures S1I and 2D). Interestingly, for the CL circuits, weaker promoters produce smaller slopes than do stronger promoters. Because transfected cells receive high plasmid copy numbers, ComMAND may operate in a resource-limited regime in transfection. In such a regime, we expect lower expression levels to generate less competition for resources, affording better control. Potentially, ComMAND performance may improve at lower DNA copy numbers than those achieved in transfection.

**Figure 2.**
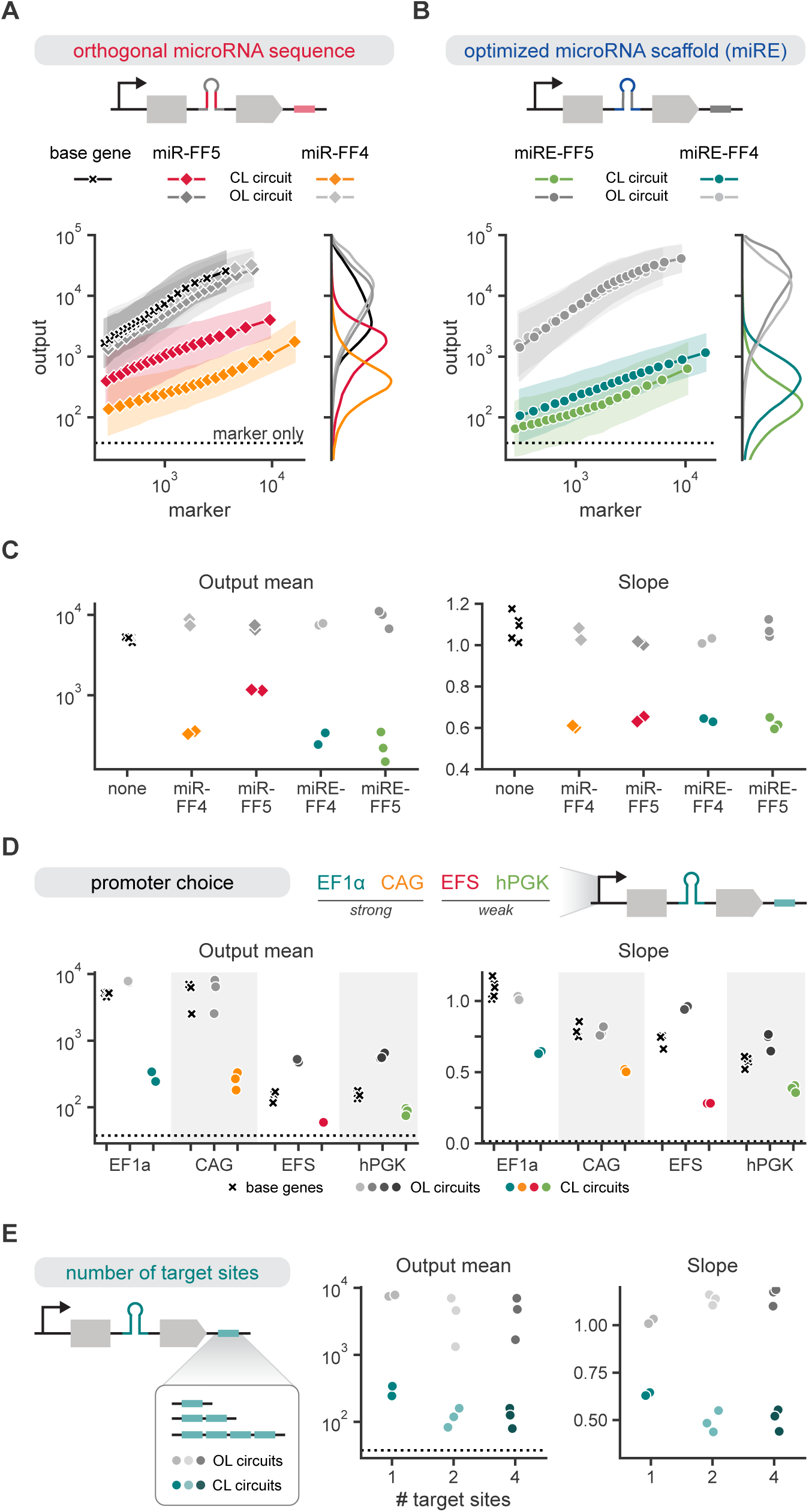
Selection of circuit components enables tuning of output expression. Three constructs with various circuit components are quantified. Base genes (x markers) include only the output gene, without intronic microRNA and target sites. Open-loop (OL, gray) and closed-loop (CL, colored) circuits combine the indicated intronic microRNA with one copy of an orthogonal (FF6, OL) or matched (CL) target site. **A**, **B**. Left: Output expression as a function of marker expression for constructs co-transfected with a marker gene in HEK293T cells. Flow cytometry measurements for one representative biological replicate are binned by marker expression into 20 equal-quantile groups per condition. Points represent geometric means of cells in each bin, and shaded regions represent this value multiplied or divided by the geometric standard deviation of the bin. Dashed line represents the output geometric mean of cells transfected only with the marker gene. Right: Histograms of output expression for cells in each condition. Circuit expression is driven from an EF1α promoter. **C**. Summary statistics of output expression for populations from 2A and 2B. **D**. Summary statistics of output expression for HEK293T cells co-transfected with a marker gene and constructs expressed from the indicated promoters. OL and CL circuits contain miRE-FF4. **E**. Summary statistics of output expression for HEK293T cells transfected with EF1α-driven miRE-FF4 circuits containing 1, 2, or 4 copies of matched (CL) or orthogonal (OL) target sites. All units are arbitrary units (AU) from a flow cytometer. Plotted mean values use the geometric mean and slope represents the slope of the line fitted to the binned, log-transformed marker-output points. Points represent individual biological replicates. Dashed lines represent values for cells transfected only with the marker gene.

Next, we explored whether changing the number of target sites could tune output expression. We first verified that changes in target site number could alter microRNA-mediated knockdown outside of an iFFL context. When we co-transfected an intronic microRNA and a separate target gene containing varying numbers of target sites, target gene expression decreased as the number of target sites increased (Figures S1F and S1G). Therefore, we expected that ComMAND would more tightly control output expression with additional target sites. However, with a second target site, the output mean and slope only slightly decreased (Figures S1J and 2E). Increasing to four target sites did not change output expression further for the CL circuit. Our observation of limited tuning via the number of target sites matches the behavior of recently reported circuits containing microRNAs paired with fully complementary target sites [30].

By varying the genetic elements in ComMAND, we were able to determine strategies to shift the output setpoint. In particular, we find that microRNA sequence, microRNA scaffold, and promoter strength can set the output expression level. Across these varied tuning strategies, we then looked to identify the underlying biochemical constraints at work in ComMAND. We turned to a mathematical model of ComMAND function to explain our experimental data, unify our understanding of the constraints, and illuminate further opportunities to modulate controller function.

### A model of ComMAND activity identifies physiological limits and tuning strategies

To achieve a better understanding of circuit properties and to more rapidly explore the wide design space, we modeled the reactions involved in ComMAND. From previously developed microRNA-based models [29], we chose a detailed set of reactions in order to account for the potentially diverse couplings of both our genetically encoded components and available cellular resources (Figure 3A, Supplemental Information Section 1). To better understand the trends we observed experimentally, we derived a steady-state analytical solution of protein output (Supplemental Information Section 1.1). Using this solution, we computed protein levels as a function of DNA copy number.

**Figure 3.**
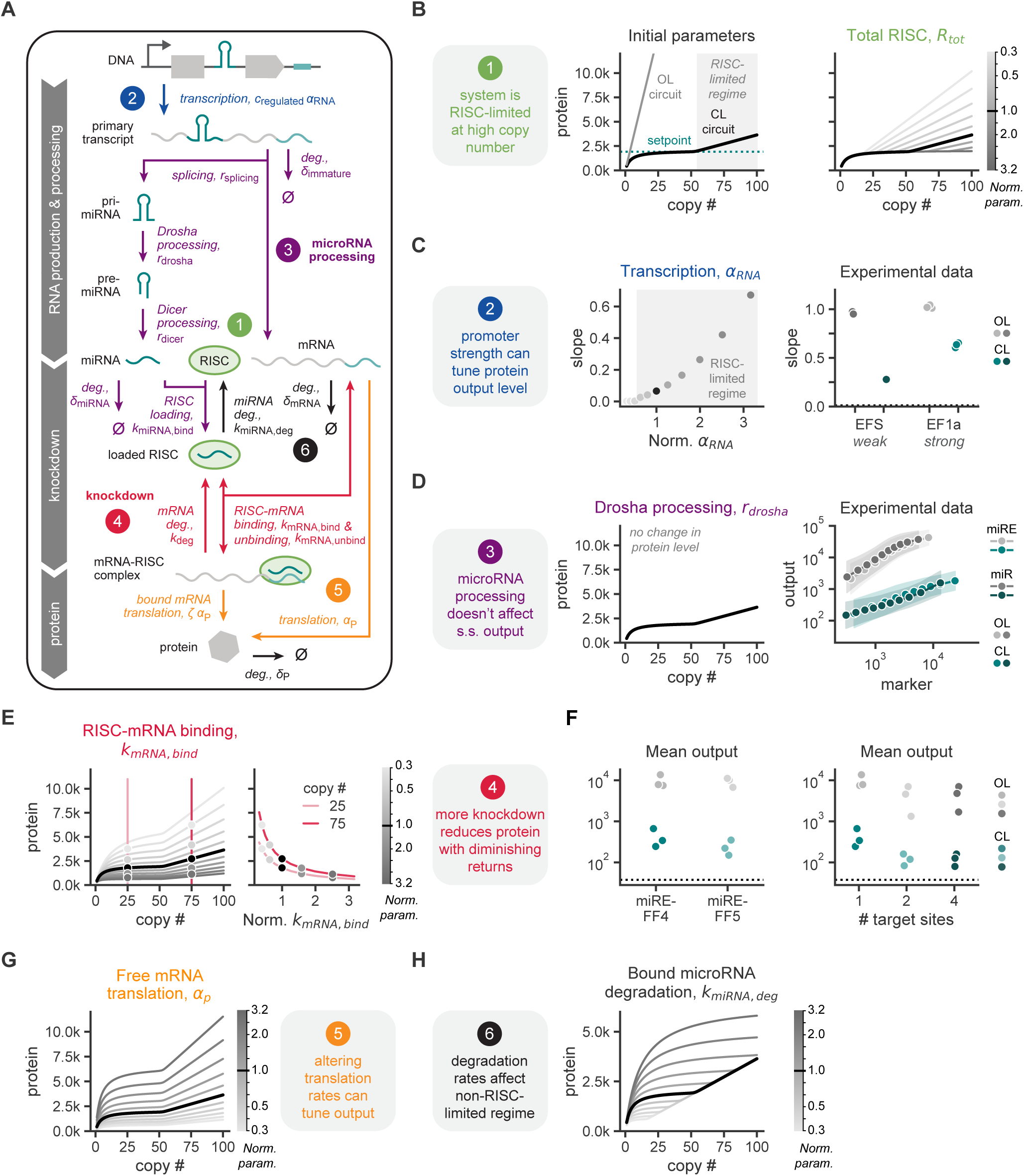
A model of ComMAND activity identifies physiological limits and tuning strategies. **A.** Schematic of a reaction network model representing circuit function. This diagram includes all species, reactions, and parameters that comprise RNA production and processing, microRNA-mediated knockdown, and protein production steps. Exact reactions, steady state analysis, and description of parameter sweeps can be found in Supplemental Information Section 1. Reactions and parameters are color-coded by design principles highlighted in the remaining panels. **B.** Left: Output protein (in molecules) plotted as a function of DNA copy number, *c*_regulated_, for the model with initial parameters as described in Supplemental Information Section 1. The curve was computed using the steady-state analytical solution. Grey shading highlights the DNA copy numbers at or above which the system becomes RISC-limited. Dashed teal line represents the system setpoint, or the protein level in the non-resource limited regime that remains constant even as copy number changes. Dashed black line indicates the solution for an unregulated gene. Right: Output protein (in thousands of molecules) as a function of DNA copy number, *c*_regulated_, for the model with a sweep of values of *R*_tot_, the total amount of RISC. Chosen parameter values (shaded grey bar) are evenly log-distributed over an order of magnitude centered on the original parameter value. The black line in this panel and subsequent panels represents the output of the model with the original parameter values, as in the left plot. **C.** Left: Slope of the copy number-protein curve at *c*_regulated_ = 100 as a function of transcription rate, *α*_RNA_. The slope of the curve is normalized by that of the unregulated gene. *α*_RNA_ values are normalized to the original value (black dot). Shaded region indicates the values of *α*_RNA_ at which the system is RISC-limited at *c*_regulated_ = 100. Right: Slope of the marker-output expression for HEK293T cells co-transfected with a marker gene and open-loop (OL, grey) or closed-loop (CL, teal) circuits expressed from EF1α or EFS promoters as in Figure 2D. Slope represents the slope of the line fitted to the binned, log-transformed marker-output points. Points represent individual biological replicates. Dashed lines represent values for cells transfected only with the marker gene. **D.** Left: Output protein (in thousands of molecules) as a function of DNA copy number for the model with a sweep of values of *r*_drosha_, the rate of Drosha processing, over one order of magnitude. All curves lie under the black line, which represents the output of the model with the original parameter values as in 3B. Right: Output expression as a function of marker expression for HEK293T cells co-transfected with a marker gene and open-loop (OL, grey) or closed-loop (CL, teal) circuits with miR-FF4 or miRE-FF4 as in Figure 2B. **E.** Left: Output protein (in thousands of molecules) as a function of DNA copy number for the model with a sweep of values of *k*_mRNA,bind_, the RISC-mRNA binding rate. Chosen parameter values (shaded grey bar) are evenly log-distributed over an order of magnitude centered on the original parameter value. Red lines highlight *c*_regulated_ = 25 and *c*_regulated_ = 75, and points indicate the values of the curves at these copy numbers. Right: Output protein (in molecules) as a function of the RISC-mRNA binding rate at two copy numbers, *c*_regulated_ = 25 and *c*_regulated_ = 75. *k*_mRNA,bind_ values are are normalized to the original value. **F.** Geometric mean output expression of constructs co-transfected with a marker gene in HEK293T cells. Left: Open-loop (OL) and closed-loop (CL) circuits with miRE-FF4 or miRE-FF5 as in Figure 2C. Right: Circuits with miRE-FF4 and varying number of target sites as in Figure S1F. Points represent individual biological replicates. Dashed lines represent geometric mean output level for cells transfected only with the marker gene. All units are arbitrary units (AU) from a flow cytometer. **G.** Output protein (in thousands of molecules) as a function of DNA copy number for the model with a sweep of values of *α_p_*, the free mRNA translation rate. Chosen parameter values (shaded grey bar) are evenly log-distributed over an order of magnitude centered on the original parameter value. **H.** Output protein (in thousands of molecules) as a function of DNA copy number for the model with a sweep of values of *k*_miRNA,deg_, the RISC-bound microRNA degradation rate,. Chosen parameter values (shaded grey bar) are evenly log-distributed over an order of magnitude centered on the original parameter value.

First, we sought to understand why output expression increased as marker expression increased, instead of remaining constant as predicted for an ideal iFFL (Figures 1D and 1E). The model predicts that protein levels remain at the circuit setpoint at lower DNA copy numbers, but exceed this setpoint at a particular copy number threshold where RISC becomes fully saturated with bound microRNA (Figure 3B, left) [29]. Above this threshold, the circuit exists in a RISC-limited regime where microRNA-mediated knockdown cannot fully control output expression. In this regime, protein levels increase linearly with copy number, and the total amount of RISC affects the rate of this increase. By changing the amount of RISC in the model, we could shift the threshold at which RISC becomes limiting as well as the slope in the RISC-limited regime (Figure 3B, right and Figure S2A, top). Because cells take up between tens and thousands of plasmids in transfection [41], our transfections may generate a RISC-limited state. This suggests that ComMAND may better regulate output expression at DNA copy numbers lower than those we obtained in transfection.

Transcriptional activity influences the rate of RISC saturation and thus the performance of ComMAND. RISC saturation is governed by the amounts of both RISC and microRNA, so we expect changes in production of primary transcript—and thus changes in microRNA levels—to shift properties of saturation. Our model predicts that promoter activity shapes the RISC-limited regime (Figure 3C, left and Figure S2B). Namely, lower transcriptional activity increases the DNA copy number threshold at which RISC becomes limiting and decreases the slope beyond this threshold. This coupling between promoter strength and RISC limitations may explain why we observe a reduced slope for the CL circuit when ComMAND is expressed from weak promoters (Figures 2D and 3C, right). The fact that we observe reduced output variability across promoters in transfection highlights that ComMAND offers control even under physiological resource-limited conditions.

Next, we explored how splicing and processing of the single primary transcript impact circuit function. Interestingly, we find that varying these parameters within one order of magnitude has little or no effect on steady state protein output (Figures S2C and 3D, left). Putatively, generation of mature microRNA from the primary transcript is not limiting at steady state for ComMAND. This may explain why the optimized miRE scaffold has no effect on output expression profiles for the FF4 circuit (Figures 2A, 2C and 3D, right).

While tuning the circuit in transfection, we hypothesized that directly increasing microRNA-mediated knockdown would enhance ComMAND performance. Indeed, in our model, increasing the binding affinity of the loaded RISC for the mature mRNA decreases the output protein setpoint and slope in the RISC-limited region (Figure 3E, left). However, this effect has diminishing returns on an absolute scale: if binding affinity is already high, further increases lead to only small decreases in protein level and slope (Figure 3E, right). This may explain why, experimentally, we find that changing the microRNA targeting sequence and the number of target sites—both potentially related to RISC-mRNA binding—has little effect on ComMAND performance (Figures 2B, 2C, 2E and 3F). Altering knockdown activity by increasing the rate of degradation of the RISC-bound mRNA demonstrates similar diminishing returns for decreasing output protein levels and slope (Figure S2D).

We next investigated how translation affects circuit output. As expected, changing the translation rate of the free mature mRNA shifts the output setpoint without changing the shape of the curve (Figure 3G). We expect that the RISC-bound mRNA can be translated, albeit at a lower rate than free mRNA. Accordingly, we assumed that the rate of bound mRNA transcription is smaller than that of free mRNA. As the ratio of translation rates between the bound and free mRNA species approaches one, the slope of the curve increases somewhat, even in the region not limited by RISC (Figure S2E). However, we observe that the “leakiness” of expression from bound mRNAs does not qualitatively change the behavior of ComMAND.

Finally, we explored how the rate of RISC-mediated degradation of the two principle species, RISC-bound microRNA and the mature mRNA, affect ComMAND function. Although we observe little impact of free microRNA degradation rate on output levels (Figure S2C), it is also possible for degradation to occur when the microRNA is loaded in RISC [42]. Interestingly, as the rate of degradation of bound microRNA increases, the output setpoint increases and the RISC saturation point shifts to higher DNA copy numbers while maintaining the same slope above the inflection point (Figure 3H). The increased control afforded by high degradation of RISC-bound microRNA suggests that a non-enzymatic mode of activity of the microRNA, where knockdown results in degradation of both the mRNA and microRNA, improves circuit performance, similar to integral controllers reported in the literature [43]. Lastly, the degradation rate of the free mRNA has only a small effect on circuit behavior (Figure S2F).

Taken together, the model helps explain our experimental results and suggests additional avenues for tuning the behavior of ComMAND. Moreover, it offers insights into circuit behavior in unconstrained and limiting resource regimes, which are largely dictated by DNA copy number, relative transcription rate, and total amount of RISC.

### The single-transcript circuit matches or exceeds performance of two-gene implementations

The single-transcript architecture of ComMAND offers the advantage of compactness; however, alternative implementations may offer additional properties. Similar iFFL circuits have been constructed using two genes, in which the microRNA and output mRNA are transcribed separately [26, 27, 30, 44]. These architectures support varying ratios of microRNA and mRNA, which can tune knockdown and may improve circuit function. To benchmark ComMAND, we compared its performance to several alternative circuit architectures (Figure 4A). In the dual-transcript implementation, the output gene containing a microRNA target site is divergently expressed with a second gene containing the intronic microRNA. In a dual-vector implementation, these genes are separately encoded on two co-delivered vectors. While we expect the dual-transcript and dual-vector implementations to have the same average expression, the dual-vector system may introduce additional extrinsic noise due to copy number variation (Figure 4A, right). When we transfected these circuits in HEK293T cells, we found that the single-transcript CL circuit controls output expression as well as or better than both two-gene architectures (Figures S3A, S3B and 4B). We observed no differences in expression of the base genes and OL circuits across all architectures, indicating that improved control is specific to circuit activity.

**Figure 4.**
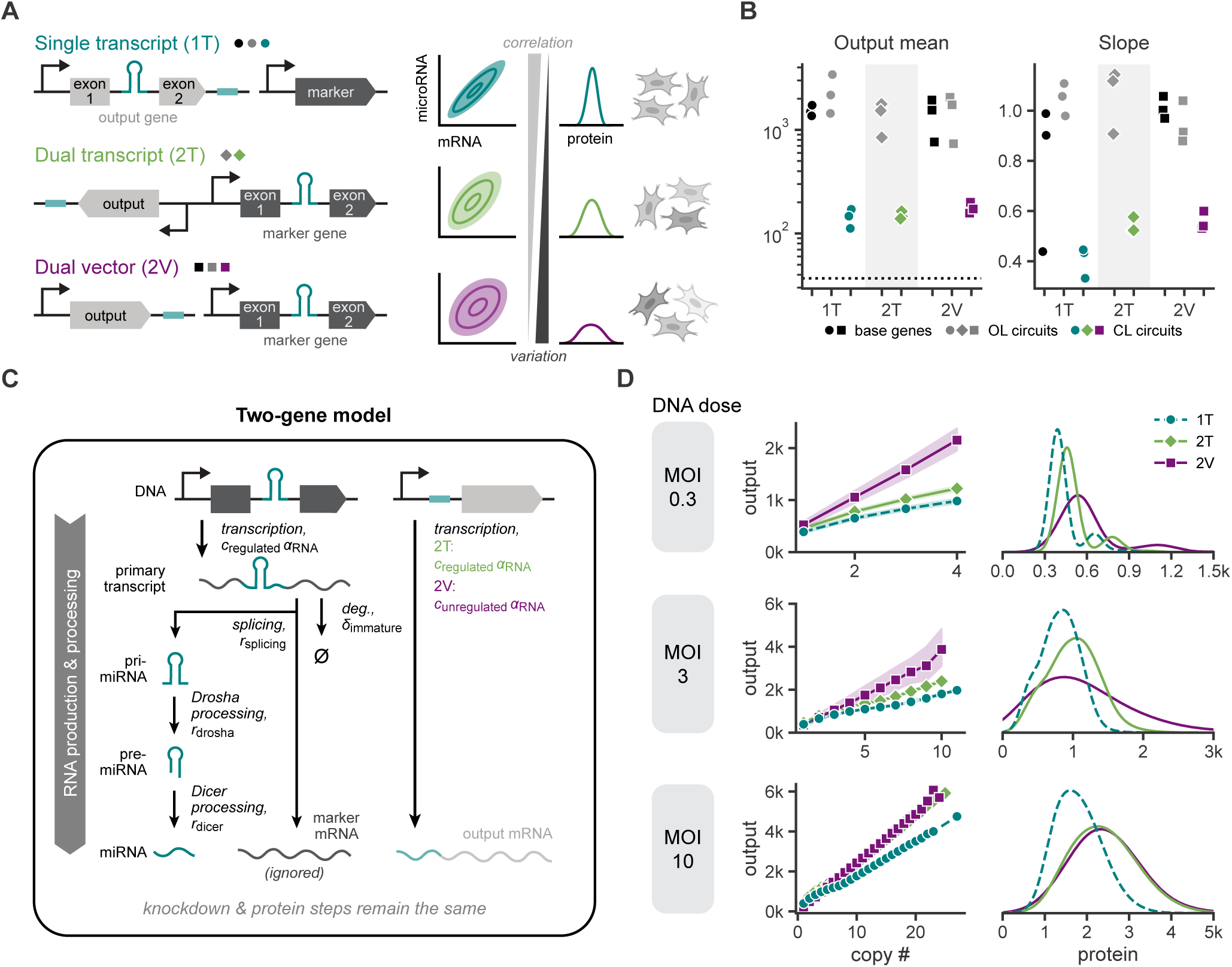
The single-transcript circuit matches or exceeds performance of two-gene implementations. **A.** Left: DNA construct diagrams of single-transcript and two-gene circuit implementations. The single-transcript circuit (ComMAND, 1T) consists of an output gene with an intronic microRNA and matched (CL) or orthogonal (OL) target sites, co-delivered with a marker gene. The dual-transcript circuit (2T) consists of a single vector with two divergently oriented genes, the marker gene with an intronic microRNA and the output gene with matched (CL) or orthogonal (OL) target sites. The dual-vector circuit (2V) consists of the same two genes as in the dual-transcript circuit, each on a separate vector. All genes are expressed from the EF1α promoter, and all circuits use miRE-FF4. Right: Delivering the circuit on a single transcript leads to strongly correlated, stoichiometrically equal expression of microRNA and mRNA species and thus to a narrow distribution of output protein expression. Dual-transcript or dual-vector architectures introduce more intrinsic noise, decreasing the correlation between microRNA and mRNA levels and increasing variation of the output protein. **B.** Summary statistics of output expression for HEK293T cells transfected with the base gene or closed-loop (CL) and open-loop (OL) miRE-FF4 circuits depicted in 4A. Presented mean values use the geometric mean. Slope represents the slope of the line fitted to the binned, log-transformed marker-output points. Points represent individual biological replicates. Dashed lines represent values for cells transfected only with a marker gene lacking an intronic microRNA. All units are arbitrary units (AU) from a flow cytometer. **C.** Schematic of RNA production and processing reactions modeled for two-gene circuit implementations. Unlike in the single-transcript model depicted in Figure 3A, here mature output mRNA is not produced by transcription and splicing of the primary transcript. Instead, it is produced directly from transcription of a second gene with transcription rate *α*_RNA_. The copy number of this second gene is either equivalent to that of the microRNA gene (*c*_regulated_, dual-transcript) or different (*c*_unregulated_, dual-vector). All other reactions remain the same as in Figure 3A. The spliced mature mRNA from the primary transcript is ignored. **D.** Left: Output protein (in thousands of molecules) as a function of DNA copy number, *c*_regulated_, from stochastic simulations of single-transcript (1T), dual-transcript (2T), and dual-vector (2V) models depicted in Figures 3A and 4C. The plot depicts summary statistics of output protein levels for the set of simulations with each copy number, where points represent the geometric mean of each set and shaded regions illustrate the geometric mean divided or multiplied by the geometric standard deviation. Right: Histograms of output protein levels (in thousands of molecules) for the same simulations. For each simulation, DNA copy numbers were chosen from a Poisson distribution with mean equal to the given MOI, the effective “multiplicity of infection” of the population, illustrated in Figure S3C. 10,000 simulations were run for each condition. See Section 1.2 for more details.

Therapeutic applications often require orders of magnitude lower DNA copy numbers than those in transfection. As copy number decreases, the effect of noise becomes more pronounced, potentially magnifying differences between circuit implementations. We expect ComMAND to outperform two-gene architectures at low DNA copy numbers because expression of circuit components are more tightly coupled. To explore circuit behavior in different copy number regimes, we turned to our model. For the two-gene architectures, we modeled output mRNA transcription separately from production of the microRNA and allowed the associated DNA copy number to vary in the dual-vector case (Figure 4C). We performed stochastic simulations using these models to account for the effects of noise, choosing DNA copy number values for each run from a distribution defined by an effective MOI (Figure S3C and Supplemental Information Section 1.2). As expected, mean output protein level increases with MOI and is higher for an unregulated gene than for the CL circuits (Figures S3D and 4D). Furthermore, output variability is lower for the single-transcript architecture than for two-gene models, especially at lower DNA copy numbers. Thus, the stochastic simulations highlight the advantage of ComMAND’s single-transcript implementation in noisy contexts.

Comparing ComMAND to two-gene, microRNA-based iFFLs, we show experimentally and via stochastic modeling that ComMAND maintains performance comparable or superior to two-gene architectures. Importantly, ComMAND maintains tight regulation across DNA copy number distributions. These results suggest that ComMAND may effectively control output expression for delivery methods with variable delivery efficiency, such as low-dosage vectors in therapeutically relevant, physiological contexts.

### Multiple designs of the single-transcript circuit regulate output expression across cell types and delivery methods

After establishing the performance of ComMAND’s single-transcript architecture, we investigated how the placement of genetic parts along the transcript affects output expression profiles. Previous work demonstrated that microRNA-mediated knockdown of mRNA is greater when target sites are located in the 5’UTR compared to in the 3’UTR [45]. Additionally, other RNA PolII-driven microRNA expression systems have placed the intronic microRNA in the 3’UTR rather than within a protein coding sequence [28, 29]. Therefore, we constructed two additional single-transcript circuit designs (Figure 5A). In the new designs, designated designs 2 and 3, we moved the microRNA target site to the 5’UTR. In design 3, we additionally moved the intronic microRNA to the 3’UTR. Transfecting these circuits in HEK293T cells, we observe that both CL circuits reduce output expression mean and variability compared to the corresponding OL circuits (Figures S4A and 5B). Output expression of both OL and CL circuits are lower for designs 2 and 3 than for the original design. Because output genes including only a microRNA or target sites also have reduced expression levels (Figures S4B and S4C), we reasoned that the location of genetic elements affects circuit activity in part by altering processes other than knockdown, such as by impeding translation of the mature mRNA (Figure 3G).

**Figure 5.**
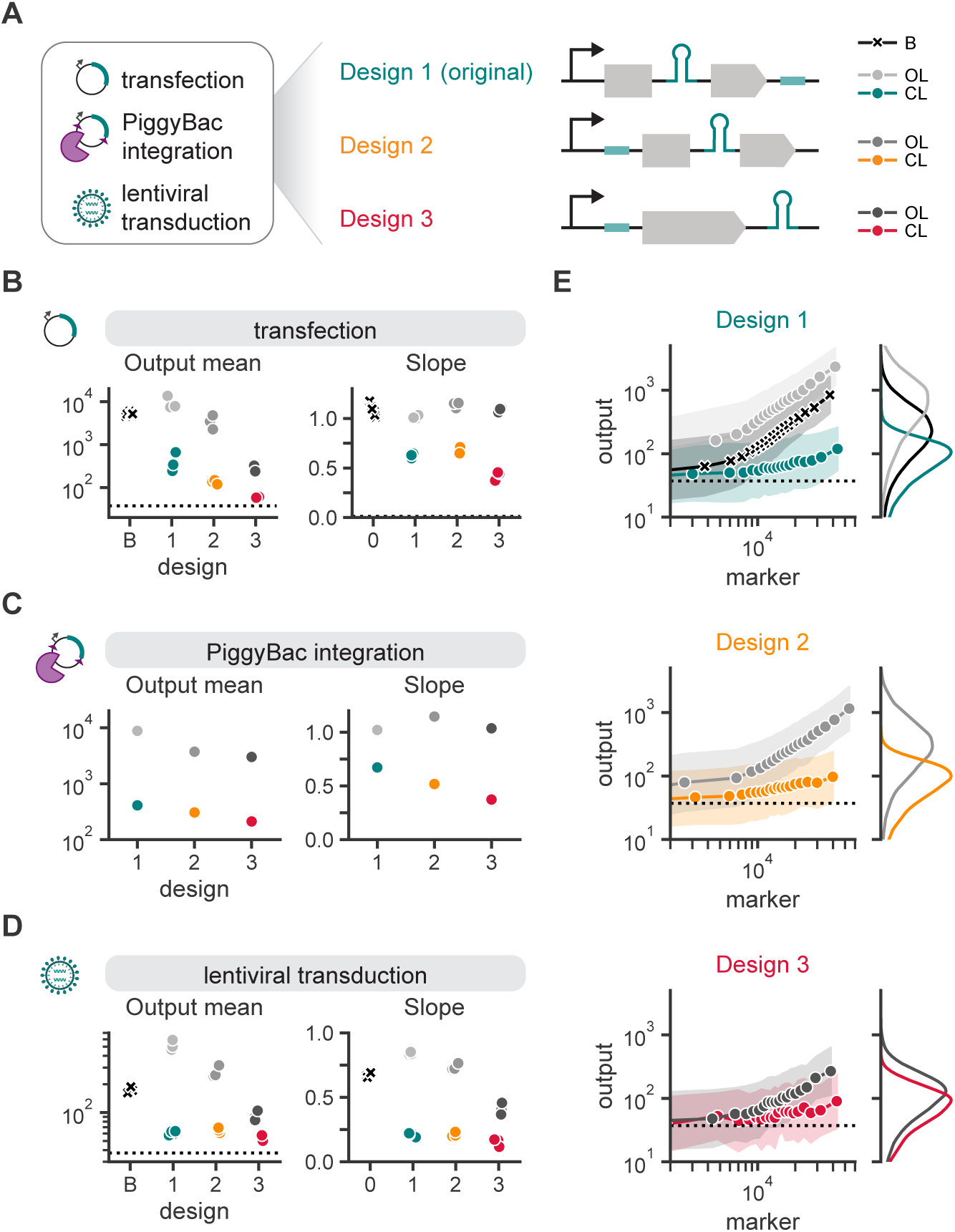
Multiple designs of the single-transcript circuit regulate output expression across cell types and delivery methods. **A.** DNA construct diagrams for three designs of the single-transcript circuit delivered to cells via transfection, PiggyBac (transposase-based) integration, and lentiviral transduction. Design 1 is the original circuit design, with an intronic microRNA within the output coding sequence and target sites in the 3’UTR. In design 2, the target site is located in the 5’UTR. In design 3, the target site is located in the 5’UTR and additionally the intronic microRNA is moved to the 3’UTR. Full vectors for PiggyBac integration and lentiviral transduction are depicted in Figure S4I and **??**, respectively. **B**, **C**, **D**. Summary statistics of output expression in HEK293T cells are shown for cells transfected (Figure 5B), PiggyBac-integrated (Figure 5C), or lentivirally transduced (Figure 5D) with constructs in Figure 5A. Plotted mean values use the geometric mean and slope represents the slope of the line fitted to the binned, log-transformed marker-output points. Points represent individual biological replicates. Dashed lines represent values for cells transfected only with the marker gene (Figure 5B) or untransduced cells (Figure 5D). Design “B” refers to the base gene construct that does not contain an intronic microRNA or target sites. **E.** Left: Output expression as a function of marker expression for one representative biological replicate of data in 5D. Points represent the geometric means of equal-quantile bins, and shaded regions represent this value multiplied or divided by the geometric standard deviation of the bin. Dashed line represents the output geometric mean of untransduced cells. Right: Histograms of output expression for cells in each condition. All units are arbitrary units (AU) from a flow cytometer.

Next, we explored whether these positional effects are element-specific or generalizable across microRNAs. To test this, we constructed circuit designs 2 and 3 with our previous panel of microRNAs. Trends in expression profiles for design 3 relative to the original design remain relatively constant across microRNAs, while those for design 2 vary across microRNAs (Figures S4D to S4F). Additionally, designs 2 and 3 perform as well as or better than corresponding two-gene architectures with target sites in the 5’UTR (Figures S4G and S4H). However, design 3 CL circuits have very low output levels, just above those for the marker-only control. Thus, the original design of ComMAND may be optimal for its consistent results across parts, though design 3 could alternatively be used to set a very low expression level. Together, these results indicate that positional and element-specific effects combine to influence output expression profiles for ComMAND.

It is possible that cell type and delivery method impact sequence-specific effects. To identify optimal designs for future therapeutic applications, we tested all three single-transcript circuit designs with miRE-FF4 in multiple expression contexts. First, we used PiggyBac transposase to randomly integrate ComMAND in HEK293T cells (Figure S4I). Again, CL circuits for all designs reduce output mean and variation compared to corresponding OL circuits (Figures S4A and 5C). Trends in mean expression are similarly preserved, where the original design has the highest output level and design 3 has the lowest. Thus, ComMAND can regulate output expression when integrated in the genome.

To expand to more application-relevant contexts, we encoded ComMAND in a lentiviral vector. To maintain high viral titer, we expressed the circuit from a small-molecule inducible promoter on the antisense strand of the virus, oriented divergently from a constitutive marker gene (Figure S5A). We transduced HEK293T cells with these viruses in the presence of inducer and found that output expression levels and variation decrease relative to unregulated genes for all three circuit designs (Figures S4A, S5B, S5C, 5D and 5E). In fact, these vectors elicit the tightest control yet for ComMAND. Viral delivery also preserves the trends in output expression between designs, further suggesting that these patterns are generalizable.

In sum, ComMAND effectively regulates output expression in transfection, transposase-based integration, and lentiviral transduction of HEK293T cells. While multiple configurations of genetic elements enable circuit function, our original design regulates output expression most consistently across contexts and thus appears appropriate for many applications. Having established the generalizability of ComMAND across delivery methods and demonstrated high performance in lentiviral vectors, we next looked to deliver the circuit to therapeutically relevant cell types.

### ComMAND functions in primary cells and can regulate clinically relevant output genes

With the ability to deliver ComMAND cargoes via lentivirus, we tested the efficacy of our gene circuit in primary cells (Figure 6A). For therapeutic applications, expression must be precisely tailored. High, unregulated expression of transgenes or heterogeneity across a population can compromise normal cell functions [16–18]. For example, overexpression of FXN in a mouse model of Friedrich’s ataxia shows limited toxicity when FXN expression remains within a range of one order of magnitude of endogenous levels but is toxic when expressed at higher levels [17, 18].

**Figure 6.**
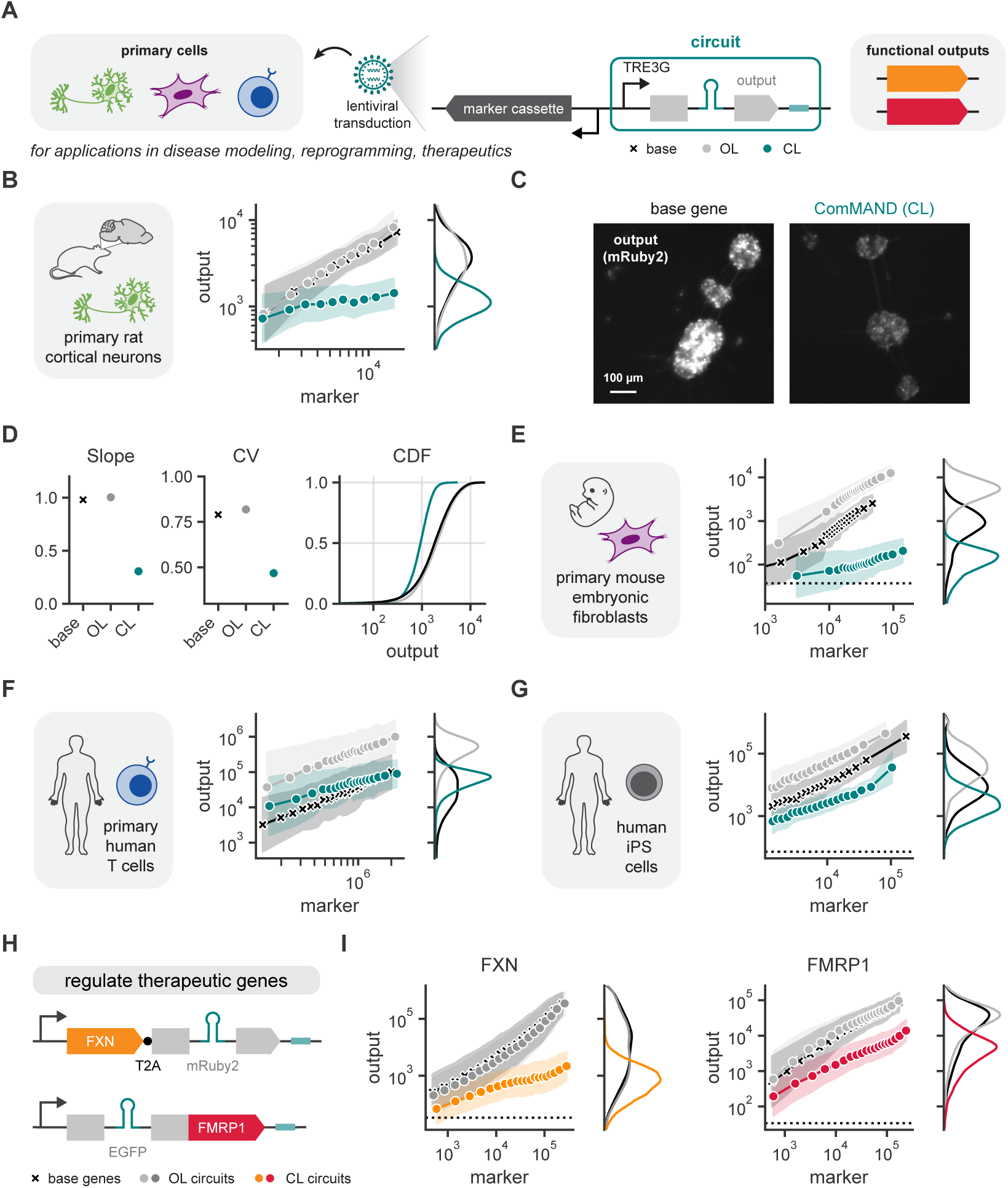
ComMAND functions in primary cells and can regulate clinically relevant output genes. **A.** Schematic of lentiviral ComMAND vectors for delivery to primary cells. The regulated gene is expressed via a doxycycline-inducible TRE3G promoter with a divergently oriented marker cassette. See Figure S5A for more details. These circuits can regulate expression of clinically relevant genes for applications in disease modeling, reprogramming, and therapeutics. **B.** Output expression as a function of marker expression for primary rat cortical neurons transduced with lentiviral vectors in Figure S5A. Data was collected in the presence of inducer (1 µg/mL doxycycline). Flow cytometry measurements are binned by marker expression into 20 equal-quantile groups per condition. Points represent geometric means of cells in each bin, and shaded regions represent this value multiplied or divided by the geometric standard deviation of the bin. **C.** Representative images of rat cortical neurons transduced with vectors expressing a base gene or the ComMAND closed-loop (CL) circuit. Images depict fluorescence of the output gene, mRuby2. **D** Summary statistics for populations presented in Figure 6B. Slope represents the slope of the line fitted to the binned, log-transformed marker-output points. CV is the coefficient of variation. The cumulative distribution function (CDF) of output expression depicts a representative biological replicate. Points represent individual biological replicates. **E** and **F**. Output expression as a function of marker expression for primary mouse embryonic fibroblasts or primary human T cells, respectively, transduced with lentiviral vectors in Figure S5A. Cells were induced and analyzed as in Figure 6B. Dashed lines represent values for untransduced cells. **G.** Output expression as a function of marker expression for human induced pluripotent stem cells co-transfected with a marker gene and base gene, open-loop circuit, or closed-loop circuit driven by the EF1α promoter. Circuits use miRE-FF4 in the original design. Cells were analyzed as in Figure 6B. Dashed lines represent values for cells transfected only with the marker gene. **H.** Schematics of ComMAND circuits used in HEK293T transfection for control of the therapeutically relevant genes FXN and FMRP1. For the FXN circuit, the microRNA is encoded in an intron witin mRuby2, which is co-expressed with FXN via a T2A peptide tag. For the FMRP1 circuit, the intronic microRNA is located within the EGFP coding sequence, and EGFP is directly fused to FMRP1. **I.** Output expression as a function of marker expression for HEK293T cells co-transfected with a marker gene and a base gene or circuit expressing fluorescently tagged frataxin (FXN, left) or FMRP1 (right). Circuits contain miRE-FF4 and are expressed via the EF1α promoter. Cells were analyzed as in Figure 6B. Dashed lines represent values for cells transfected only with the marker gene.

We investigated the performance of ComMAND circuits by transducing the base gene, OL circuit, and CL circuit in three primary cell types: rat cortical neurons (Figures 6B to 6D), mouse embryonic fibroblasts (Figure 6E), and human T cells (Figure 6F). T cells are used for a variety of immunotherapies, whereas mouse embryonic fibroblasts and rat cortical neurons are important model cell types for cellular reprogramming and neurological gene therapy delivery, respectively. In all of these cell types, ComMAND successfully reduces mean expression and narrows the expression distribution of a model fluorescent protein cargo (Figures S5D to S5J and 6B to 6F).

For the rat cortical neurons, we quantified the slope of the output distribution and observed strongly sublinear scaling with respect to marker expression, indicating tight control. We find that ComMAND dramatically reduces population heterogeneity as measured by the coefficient of variation—graphically visible in the cumulative distribution function as a steeper slope—relative to the unregulated controls (Figure 6D). Observing clusters of transduced neurons with microscopy (Figures S5E and 6C), we find that ComMAND generates more uniform expression across and between clusters of neurons.

We also tested ComMAND activity in transfection of human induced pluripotent stem cells (iPSCs). iPSCs are increasingly used as a starting cell type for disease modeling and to produce diverse cell therapies [19, 46, 47]. Similarly, we find that ComMAND reduces output protein mean and slope in iPSCs (Figures S6A to S6C and 6G).

With the ability to control transgenes in therapeutically relevant delivery contexts and cell types, we sought to use ComMAND to regulate genes affected in two monogenic neurological disorders that have a narrow window of therapeutic efficacy. Fragile X results from loss of expression of FMRP1 and is the most common inherited form of mental retardation [48].

Friedrich’s ataxia is a progressive neurodegenerative disease that results from the loss of frataxin (FXN) [49]. To model gene therapies for Fragile X and Friedrich’s ataxia, we sought to control expression of FMRP1 and FXN in transfection of HEK293T cells as a proof of concept. We expressed FXN alongside a fluorescent protein encoding the microRNA-containing intron using a self-cleaving 2A peptide sequence (Figure 6H, top). For FMRP1, we added the microRNA-containing intron to a fluorescent protein directly fused to the gene (Figure 6H, bottom). In both of these cases, ComMAND reduces the mean and slope of the output protein expression, increasing the proportion of cells operating under a theoretical safe limit of expression (Figures S6D, S6E and 6I). This proof-of-concept work lays the foundation for translation of ComMAND to therapeutically relevant contexts where tight control of transgene expression is essential for safety and efficacy.

## Discussion

To build gene circuits that provide controlled dosages of transgenic cargoes, we developed ComMAND, a single-transcript, microRNA-mediated incoherent feedforward loop (Figure 1). As an RNA-based control system, ComMAND provides compact and modular control, minimal immunogenicity, low resource burden, orthogonality to native gene networks, and programmability via selection of sequences [50]. ComMAND achieves sublinear scaling of transgene dosage, approaching the theoretical limit of control at low, clinically relevant DNA doses (Figure 2). As ComMAND operates at the level of post-transcriptional regulation, the dosage of the transgenic cargo delivered by ComMAND can be orthogonally tuned via promoter selection. Following rational tuning of components, we used mathematical modeling to examine constraints on ComMAND performance and to identify design rules (Figure 3). Further, we used stochastic modeling coupled with experiments to compare ComMAND to alternative two-gene architectures and identify optimal regimes of performance (Figure 4). In delivering diverse transgenic cargoes, we demonstrate that ComMAND tightly controls expression across diverse cell types including human induced pluripotent stem cells, primary mouse embryonic cells, primary rat neurons, and primary human T cells (Figures 5 and 6). Importantly, we show that ComMAND constrains expression of the therapeutically relevant transgenes FXN and FMRP1 within a narrow window, supporting translational therapies for two neurological disorders.

Transgene dosage from ComMAND can be rationally tuned by selection of genetic parts. Stronger promoters increase expression of transgenes from ComMAND, and optimization of the microRNA sequence and scaffold can improve mean expression and reduce variability, albeit with diminishing returns (Figures 2A to 2C). While addition of multiple target sites tunes the output in other microRNA-mediated iFFLs, we observe only a small effect from increasing the number of target sites (Figure 2E). However, our target sites are perfectly complementary to the microRNA. Because perfectly and imperfectly complementary sequences are processed via different mechanisms, ComMAND may have a relatively small range of tunability with respect to target site number [1, 30, 33, 44]. As previously observed for other iFFLs [28, 43], ComMAND reduces resource burden, extending the range of co-expression for the marker and the output compared to the base gene (Figure 2A).

We used our mathematical model to understand the parameters that physiologically constrain the performance of ComMAND. We identify the ratio of the transcription rate and the total pool of RISC as a sensitive parameter that defines the regime of controllability for ComMAND (Figures 3B and 3C). Specifically, saturation of RISC sets a physiological limit on the control afforded by microRNA-mediated circuits. As total levels of RISC are regulated through a myriad of loading and degradation mechanisms that can vary by cell type and genetic background [51, 52], it is more feasible to tune ComMAND to reside in the regime of controllability through promoter selection. We also identify that most parameters affecting microRNA processing do not significantly impact output expression, suggesting that optimization of these steps may not be fruitful (Figures S2C and 3D).

We designed ComMAND as a single transcript for optimal compactness and tight coupling between the production of mRNA and microRNA species. In theory, a single-transcript, microRNA-mediated iFFL provides optimal noise suppression [53]. Dosage control from ComMAND matches or exceeds the performance of dual-transcript and dual-vector architectures in transfection of HEK239T cells (Figure 4B). Transfections deliver hundreds of copies of plasmids, potentially obscuring differences in designs that would manifest at lower copy number. Examining a range of low copy numbers with stochastic simulations, we find that ComMAND outperforms two-gene implementations in this clinically relevant regime (Figure 4D). Thus, we conclude that single-transcript designs will provide optimal control of therapeutic cargoes.

While ComMAND provides precise control of transgene expression by mitigating variance inherent to methods of delivery, there remain limits to this control and opportunities to further define design parameters. As a form of negative regulation, we expect that ComMAND will decrease expression of the output gene. In general, we observe reductions in the mean expression relative to the open-loop circuit and base gene. However, introduction of the intronic microRNA increases expression from ComMAND compared to the base gene in T cells (Figure S5I). Addition of an intron may increase expression through splicing-mediated transcriptional activation and may be differentially regulated across cell types [54, 55]. As the position of introns influences transcription, there remains an opportunity to understand how placement of the intronic microRNA along ComMAND’s single transcript tunes expression for therapeutically relevant genes and varies across cell types. In theory, ComMAND controls for DNA copy number as well as transcriptional variance to reduce noise in gene expression [53]. However, significant variance introduced in translation may obscure noise suppression at the transcriptional level. Other forms of regulation such as translational control and protein-based circuits may be coupled with ComMAND to control expression across the Central Dogma [56–59].

Here, we show that ComMAND controls transgene dosage in a range of mammalian primary cells including primary human T cells and iPSCs, demonstrating the enormous potential to apply ComMAND to control transgenes for therapeutic applications and basic research. For haploinsufficiency disorders such as Friedrich’s ataxia and Fragile X, ComMAND-regulated gene supplementation of FXN and FMRP1 may offer safe, effective gene therapies. As a post-transcriptional regulator, ComMAND can putatively control cargoes from cell type-specific promoters for targeted expression. Additionally, integration of cell-state and pathway-responsive promoters may scale expression of ComMAND for cell-autonomous feedback. As transgenes are increasingly used to augment cellular functions and program cell fate, ComMAND may support well-controlled expression of transgenes for diverse therapeutic applications [60–62]. By increasing the predictability of transgene expression, we expect ComMAND will broadly improve the performance of gene circuits in therapeutic contexts.

## Acknowledgements

Research reported in this manuscript was supported by the National Institute of General Medical Sciences of the National Institutes of Health under award number R35-GM143033, by the National Science Foundation with the NSF-CAREER under award number 2339986, and the with funding from the Institute for Collaborative Biotechnologies. K.S.L. and E.L.P. are supported by the National Science Foundation Graduate Research Fellowship Program under grant number 1745302. We thank Jack Toppen, Albert Blanch Asensio, Adam Beitz, Deon Ploessl, and Diya Godavarti for help with experiments. We thank Sneha Kabaria, Nathan Wang, Deon Pleossl, and Yunbeen Bae for feedback on the development of the manuscript. iPS11 cells were a gift from the Weiss lab, and the PiggyBac supertransposase plasmid was a gift from the Elowitz lab.

## Author Contributions

K.S.L. and K.E.G. conceived and outlined the project. K.S.L., C.P.J., E.L.P., and S.G. performed experiments. K.S.L. and C.P.J. analyzed experiments. C.P.J. and E.L.P. composed the model. K.S.L., K.E.G., C.P.J., and E.L.P. wrote and edited the manuscript. K.E.G. supervised the project.

## Data and Materials Availability

Raw data available upon request. Code for data analysis and modeling will be made available at https://github.com/GallowayLabMIT/.

## Methods

### Cloning

The sequences for FF3, FF4, FF5, and FF6 microRNA hairpins and cognate target sites were obtained from Leisner *et al.* [39]. To clone the microRNAs, oligonucleotides were ordered from Azenta/Genewiz, phosphorylated, annealed, and ligated into a microRNA scaffold digested by XhoI and EcoRI. The miR-30a scaffold was PCR amplified from Addgene vector #25748, and the miR-E scaffold was ordered as a gBlock from Azenta/Genewiz based on the sequence in Nissim *et al.* [40]. The intronic microRNAs were then introduced into mRuby2, mGreenLantern, EGFP, and mCherry coding sequences at an AGGT site via Gibson assembly using Hifi DNA Assembly Master Mix (NEB, M5520) according to manufacturer’s instructions.

To clone the microRNA target sites, oligonucleotides were ordered from Azenta/Genewiz, phosphorylated, annealed, and ligated into a “part vector” backbone digested by HindIII and AvrII.

The therapeutically relevant genes, FMRP and FXN, were ordered from Addgene (#87929 and #23620) and cloned into “part vectors” via Gibson assembly using Hifi DNA Assembly Master Mix according to manufacturer’s instructions.

All plasmids for transfection were constructed via BsaI (NEB, R3733L) Golden Gate cloning using “part vectors” corresponding to promoter, coding sequence, UTR, and polyadenylation signal. These single transcriptional units were then combined with a backbone vector and a 200-bp spacer via PaqCI (NEB, R0745L) Golden Gate cloning to construct the lentiviral and PiggyBac integration vectors. The lentiviral backbone is a third-generation vector derived from Addgene #17297.

### HEK293T cell culture and transfection

HEK293T cells were cultured using DMEM (Genesee Scientific, 25-501) plus 10% FBS (Genesee Scientific, 25-514H) with 0.1% gelatin coating (Sigma-Aldrich, G1890-100G) and incubated at 37*^◦^*C with 5% CO_2_. HEK293T cells were dissociated using 0.25% Trypsin-EDTA (Genesee Scientific, 25-510) diluted in PBS (Sigma-Aldrich, P4417-100TAB) for four minutes then quenched with an equal volume of DMEM + 10% FBS.

For experiments, cells were counted using a hemocytometer and plated at a density of 25,000-35,000 cells per 96-well 24 hours before transfection. Transfection was performed using linear polyethylenimine, PEI (Fisher Scientific, 4389603). Transfection mixes were prepared using a ratio of 4 µg PEI to 1 µg DNA. First, a master mix of PEI and KnockOut™ DMEM (ThermoFisher Scientific, 10-829-018) was prepared and incubated for a minimum of 10 minutes. This mixture was then added to DNA mixes containing 112.5 ng of output plasmid and 56.25 ng of marker plasmid per well. These conditions mixes were incubated further for 10 to 15 minutes and then added on top of the growth media in the 96-well plate. After 24 hours, media was replaced with fresh DMEM + 10% FBS. At 2 days post transfection, cells were prepared for flow cytometry by disassociating using 0.25% Trypsin-EDTA diluted in PBS for four minutes then quenching with an equal volume of DMEM + 10% FBS. After centrifuging at 500x*g* for 5 minutes, cells were resuspended in PBS and transferred to a v-bottom plate for flow cytometry.

### iPSC culture and transfection

iPS11 cells (Alstem) were cultured using mTeSR™ Plus (STEMCELL Technologies, 100-1130) with Geltrex™ (ThermoFisher Scientific, A1413302) coating and incubated at 37*^◦^*C with 5% CO_2_. For passaging, iPS11 cells were dissociated in clumps using ReLeSR™ (STEMCELL Technologies, 100-0484) according to the manufacturer’s instructions. For experiments, iPS11 cells were dissociated as single cells using Gentle Cell Dissociation Reagent (STEMCELL Technologies, 100-1077) according to manufacturer’s instructions and counted using a hemocytometer. Cells were plated in mTeSR™ Plus with 10 µM ROCK inhibitor (Millipore Signma, Y0503-5MG) and 100 U/mL penicillin-streptomycin (Gibco, 15140122).

For experiments, 15,000 iPS11 cells were plated per well in 96-wells 48 hours before transfection. 24 hours before transfection, media containing ROCK inhibitor was removed and replaced with mTeSR™ Plus with penicillin-streptomycin. On the day of transfection, the media was changed to Opti-MEM™ (ThermoFisher Scientific, 31985062) and transfection mixes were prepared with FUGENE® HD (FuGENE, HD-1000) according to the manufacturer’s instructions using a ratio of 3 µL reagent to 1 µg DNA. Each well was transfected with 66 ng output plasmid and 33 ng marker plasmid. 4 hours after transfection, mTeSR™ Plus with penicillin-streptomycin was added to the wells. 24 hours after transfection, the media was changed to mTeSR™ Plus with penicillin-streptomycin. 2 days post transfection, cells were detached from wells using Gentle Cell Dissocation Reagent and centrifuged at 500x*g* for 5 minutes. Cells were resuspended in PBS and transferred to a v-bottom plate for flow cytometry.

### PiggyBac integration

For PiggyBac integration, 800,000 HEK293T cells were plated per well of a 6-well plate coated with 0.1% gelatin. The following day, 1.8 µg of the integration vector and 600 ng of a plasmid expressing the PiggyBac supertransposase (gift from the Elowitz lab) per well were co-transfected using PEI. Specifically, a master mix was created by adding 9.6 µg per well of PEI to 269 µL per well of KnockOut™ DMEM (ThermoFisher Scientific, 10-829-018) and incubated at room temperature for 15 minutes. Then, this master mix was added to the 2.4 µg of plasmid DNA per condition and allowed to incubate at room temperature for 15 minutes. The resulting condition mixes were added drop-wise to the wells.

The day after transfection, media was replaced with DMEM + 10% FBS plus 1 µg/mL puromycin (InvivoGen, ant-pr-1). This was repeated for four more days. On the sixth day post-transfection, cells were prepared for flow cytometry by dissociating with 0.25% Trypsin-EDTA diluted in PBS for four minutes followed by quenching with an equal volume of DMEM + 10% FBS. After centrifuging at 500x*g* for 5 minutes, cells were resuspended in PBS and transferred to a v-bottom plate for flow cytometry.

### Lentivirus production

Three separate passages of Lenti-X HEK293T cells (Takara Bio, 632180) grown in DMEM + 10% FBS were seeded at 10^6^ cells per well of a 6-well plate. The following day (day 1), 1 µg of the third-generation lentiviral expression plasmid, 1 µg of the packaging plasmid (psPAX2, Addgene #12260), and 2 µg of the envelope plasmid (pMD2.G / VSVG, Addgene #12259) per well were co-transfected using PEI. Specifically, a master mix was created by adding 16 µg per well of PEI to 222 µL per well of KnockOut™ DMEM and incubated at room temperature for 15 minutes. Then, this master mix was added to the 4 µg of plasmid DNA per condition and allowed to incubate at room temperature for 15 minutes. The resulting condition mixes were added drop-wise to the wells. After 6 hours, the media was aspirated and replaced with 1.25 mL of DMEM + 10% FBS + 25 mM HEPES (Sigma-Aldrich, H3375). On the following day (day 2), the media was collected, stored at 4*^◦^*C, and replaced with HEPES-buffered DMEM + 10% FBS. On day 3, the media was again collected. The collected media was filtered through a 0.45 µm PES filter.

To the filtered virus-containing media, Lenti-X Concentrator (Takara Bio, 631232) was added in a 3 parts media : 1 part concentrator volume ratio, mixed gently, and left overnight at 4*^◦^*C. On day 4, the media was centrifuged at 1500x*g* at 4*^◦^*C for 45 minutes. The supernatant was aspirated, and the resulting pellet was resuspended to a total volume of 200 µL in HEPES-buffered DMEM + 10% FBS. Virus was used immediately or stored at -80*^◦^*C.

For the T cell and rat cortical neuron experiments, virus was produced as above at 10-cm dish scale, using 6 µg/dish of the lentiviral expression plasmid, 6 µg/dish of the packaging plasmid, and 12 µg/dish of the envelope plasmid. 6.5 mL of HEPES-buffered DMEM + 10% FBS per dish was collected on successive days, and the final pellet was resuspended to a total volume of 500 µL.

### HEK293T lentiviral transduction

On the day of transduction, HEK293T cells were dissociated using 0.25% Trypsin-EDTA diluted in PBS, and cells were counted using a hemocytometer and diluted to a concentration of 20,000 cells per well in DMEM + 10% FBS. Cells were combined with 5 µg/mL polybrene (hexadimethrine bromide, Sigma-Aldrich, H9268-5G) and either a constant amount of virus (1.0 µL per well) or a two-fold serial dilution of the produced virus (highest concentration: 1.0 µL concentrated virus per well). Additional DMEM + 10% FBS was added for a total volume of 100 µL per well. The resulting cell, polybrene, and virus mixture was plated onto 96-well plates coated with 0.1% gelatin.

Six hours later, the cells were media changed into fresh DMEM + 10% FBS containing 1 µg/mL doxycycline (dox, Sigma-Aldrich, D3447-500MG). The following day, the media was replaced with fresh dox-containing media. Cells were prepared for flow cytometry three days later (4 days post-transduction) by disassociating using 0.25% Trypsin-EDTA diluted in PBS for four minutes then quenching with an equal volume of DMEM + 10% FBS. After centrifuging at 500x*g* for 5 minutes, cells were resuspended in PBS and transferred to a v-bottom plate for flow cytometry.

### Primary mouse embryonic fibroblast lentiviral transduction

Primary mouse embryonic fibroblasts were isolated as described in Wang *et al.* [63]. Vials of cryopreserved passage 1 cells were thawed into DMEM + 10% FBS in T75 flasks coated with 0.1% gelatin. After 1-2 days of recovery, cells were dissociated using 0.25% Trypsin-EDTA diluted in PBS. Cells were counted using a hemocytometer and plated onto 96-well plates coated with 0.1% gelatin at 10,000 cells per well.

The following day, the media was replaced with DMEM + 10% FBS plus 5 µg/mL polybrene and either a constant amount of virus (10.0 µL/mL) or a two-fold serial dilution of virus (highest concentration: 20 µL concentrated virus per well). The cells were spinfected by centrifuging at 1500x*g* for 90 minutes at 32*^◦^*C.

Six hours after the spinfection, the cells were changed into fresh DMEM + 10% FBS containing 1 µg/mL doxycycline. The cells were changed to fresh dox-containing media the following day. Cells were prepared for flow cytometry three days later (4 days post-transduction) by disassociating using 0.25% Trypsin-EDTA diluted in PBS for four minutes then quenching with an equal volume of DMEM + 10% FBS. After centrifuging at 500x*g* for 5 minutes, cells were resuspended in PBS and transferred to a v-bottom plate for flow cytometry.

### Primary rat cortical neuron lentiviral transduction

Rat cortical neurons (Thermo Fisher, A36511) were recovered following manufacturer’s instructions. Neurons were cultured in Neurobasal Medium (Thermo Scientific, 21103049) supplemented with 0.5 mM GlutaMAX (Thermo Fisher Scientific, 35050061) and 20 mL/L B-27 (Thermo Scientific, 17504044). A 48-well plate was coated first with 0.1% gelatin followed by 10 mg/mL laminin (Corning, 354232). Neurons were then plated at 80,000 cells per well. Six hours after plating, a half-media change was performed, where half the media volume was aspirated and replaced with fresh pre-warmed media. After this point, a half-media change with pre-warmed media was performed every day.

Two days after plating, the cells were transduced with virus added at a multiplicity of infection (MOI) of 1, 5, or 7 with 5 µg/mL of polybrene. MOI was estimated from viral titers computed from the mouse embryonic fibroblast transduction. Six hours after transduction, the cells were half-media changed twice into supplemented Neurobasal media prepared as above with the addition of 1 µg/mL doxycycline. This double media change was performed again on the two subsequent days. Three days after transduction, the cells were imaged and prepared for flow cytometry. The neurons were disassociated gently using DMEM/F12 (Corning, 10-090-CV) with 17 U/mL DNase (Worthington Biochemical, LK003172) and 167 U/mL papain (Worthington Biochemical, LK003178) at 37*^◦^*C for 40 minutes. The resulting clusters of neurons were centrifuged at 400x*g* for 4 minutes, resuspended in PBS, and transferred to a v-bottom plate for flow cytometry.

### Primary human T cell lentiviral transduction

Peripheral blood mononuclear cells from a healthy donor were purified from a leukopak (STEMCELL Technologies, 200-0092) using a EasySep Direct Human PBMC Isolation Kit (STEMCELL Technologies, 19654) according to manufacturer’s instructions. Primary CD8+ T cells were isolated using EasySep Human CD8+ T Cell Enrichment kits (STEMCELL Technologies, 19053) and cultured in RPMI-1640 (ATCC, 30-2001) supplemented with 10% FBS, penicillin-streptomycin, and 30 IU/mL recombinant human IL-2 (R&D Systems). Following isolation, cells were activated with DynaBeads Human T-Activator CD3/CD28 (Thermo Fisher, 11131D) at a 1:1 cell-to-bead ratio and cultured in 24-well plates with 1 million cells per mL at 37*^◦^*C and 5% CO_2_ overnight.

For transduction the following day, cells were re-plated at 1 million cells per mL (500,000 cells per condition), and concentrated lentiviruses were added to cells at an MOI of 1, 5, or 7 with 8 µg/mL diethylaminoethyl-dextran (Sigma-Aldrich, D9885). MOI was estimated from viral titers computed from the mouse embryonic fibroblast transduction. On day 4 post-isolation, DynaBeads were removed according to manufacturer’s instructions and cells were expanded in fresh IL-2-containing media. On day 7, cells were treated with 1 ug/mL doxycycline. After 24 hours, cells were stained with LIVE/DEAD fixable near-IR dye (ThermoFisher, L34975) according to manufacturer’s instructions and analyzed using a Cytoflex S flow cytometer.

### Flow cytometry

For all experiments except those with the primary human T cells, flow was performed with an Attune NxT flow cytometer. See Supplemental Information Table S1 for channel mappings and voltages. Data were analyzed by selecting single cells using forward and side scatter gates. Cells were then gated on marker expression based on the 99.9^th^ percentile of the untransfected or untransduced populations.

The primary human T cells were analyzed using a Cytoflex S flow cytometer. See Supplemental Information Table S2 for channel mappings and voltages. Data were analyzed by selecting single cells using forward and side scatter gates. Live cells were gated as the negative population of the live/dead dye, and *>*96% of cells were viable. Cells were then gated on marker expression based on the 99.995^th^ percentile of the untransduced population.

