## Supplementary Figures for "Model-guided design of microRNA-based gene circuits supports precise dosage of transgenic cargoes into diverse primary cells"

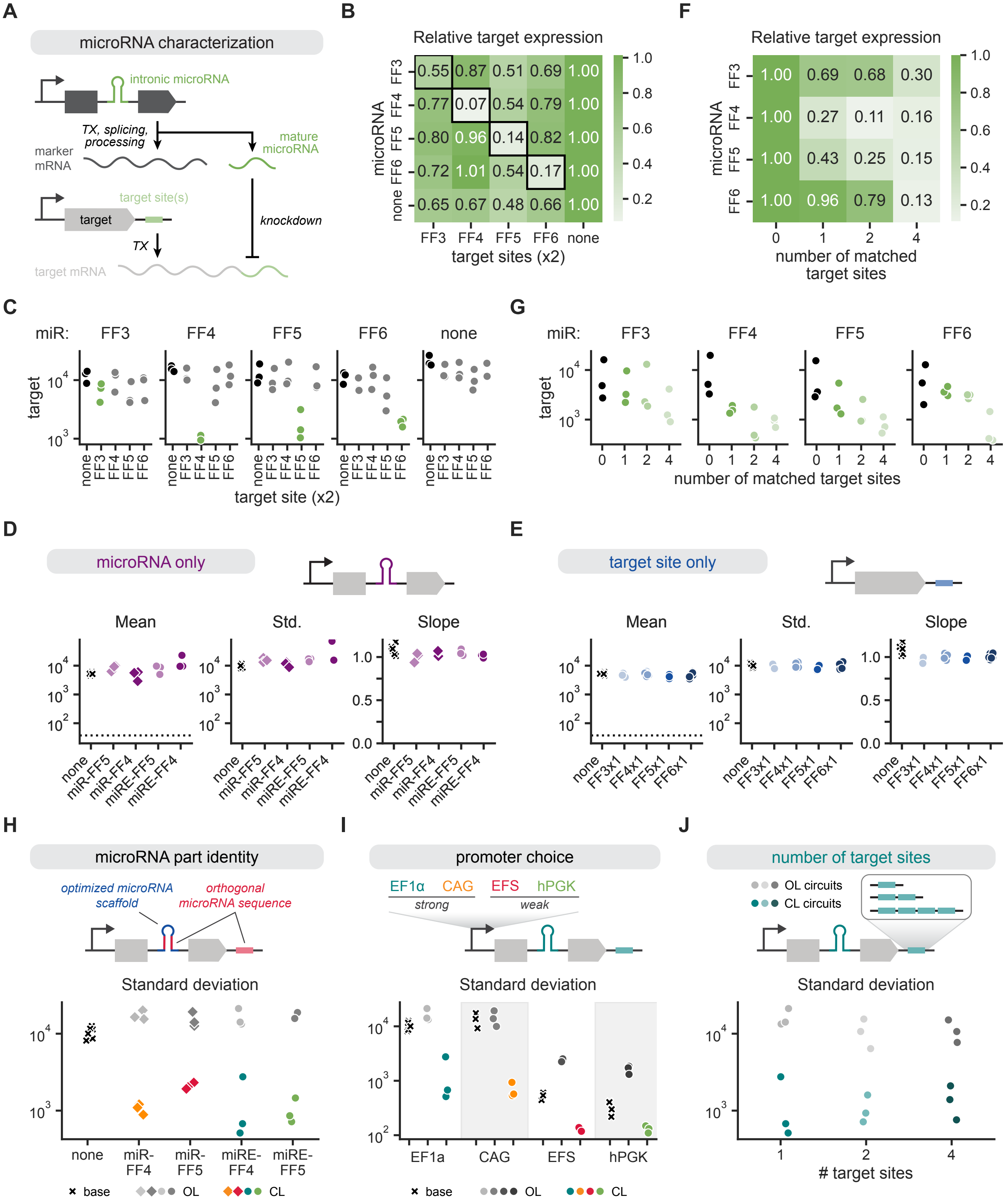

**Figure S1. Separate microRNA and target site characterization and additional circuit tuning data.**

**A**. Schematic of constructs for microRNA and target site characterization. A construct with a miR-30a–based intronic microRNA (miR-FF3, miR-FF4, miR-FF5, or miR-FF6) is transcribed, spliced, and processed into mature marker mRNA and mature microRNA. A co-delivered construct containing a target gene with 1, 2, or 4 target sites (FF3, FF4, FF5, or FF6 target sequences) is transcribed into mature target mRNA, which can be knocked down by the cognate microRNA.

**B**. Heatmap of relative target gene expression for microRNA and target site sequence combinations in HEK293T cells co-transfected with constructs in Figure S1A. Each marker-microRNA construct was paired with a target gene containing two copies of the indicated target site sequence. Population geometric means were calculated for each condition, and expression levels were then normalized for each microRNA (rows) to the no target site condition (rightmost column) for each biological replicate. Final values represent averages of $n = 3$ biological replicates. Black outlines indicate conditions with matched microRNA and target site sequences.

**C**. Geometric mean of target gene expression for conditions summarized in Figure S1B.

**D**, **E**. Summary statistics for HEK293T cells transfected with output genes containing only an intronic microRNA (S1D) or only a 3'UTR target site (S1E). All constructs were expressed by an EF1α promoter. The plotted mean values use the geometric mean. Std. refers to the standard deviation, and the slope represents the slope of the line fitted to the binned, log-transformed marker-output points. Dashed lines represent values for cells transfected only with the marker gene.

**F**. Heatmap of relative target gene expression for microRNA and target site number combinations in HEK293T cells co-transfected with constructs in S1A. Each marker-microRNA construct was paired with a target gene containing 0, 1, 2, or 4 copies of the matched target site sequence. Population geometric means were calculated for each condition, and expression levels were then normalized for each microRNA (rows) to the no target site condition (leftmost column) for each biological replicate. Final values represent averages of $n = 3$ biological replicates.

**G**. Geometric mean of target gene expression for conditions summarized in Figure S1F.

**H**. Output standard deviations for populations from Figure 2A and Figure 2B.

**I**. Output standard deviations for populations from Figure 2D.

**J**. Output standard deviations for populations from Figure 2E.

Points represent individual biological replicates. All unnormalized values are in arbitrary units (AU) from a flow cytometer.

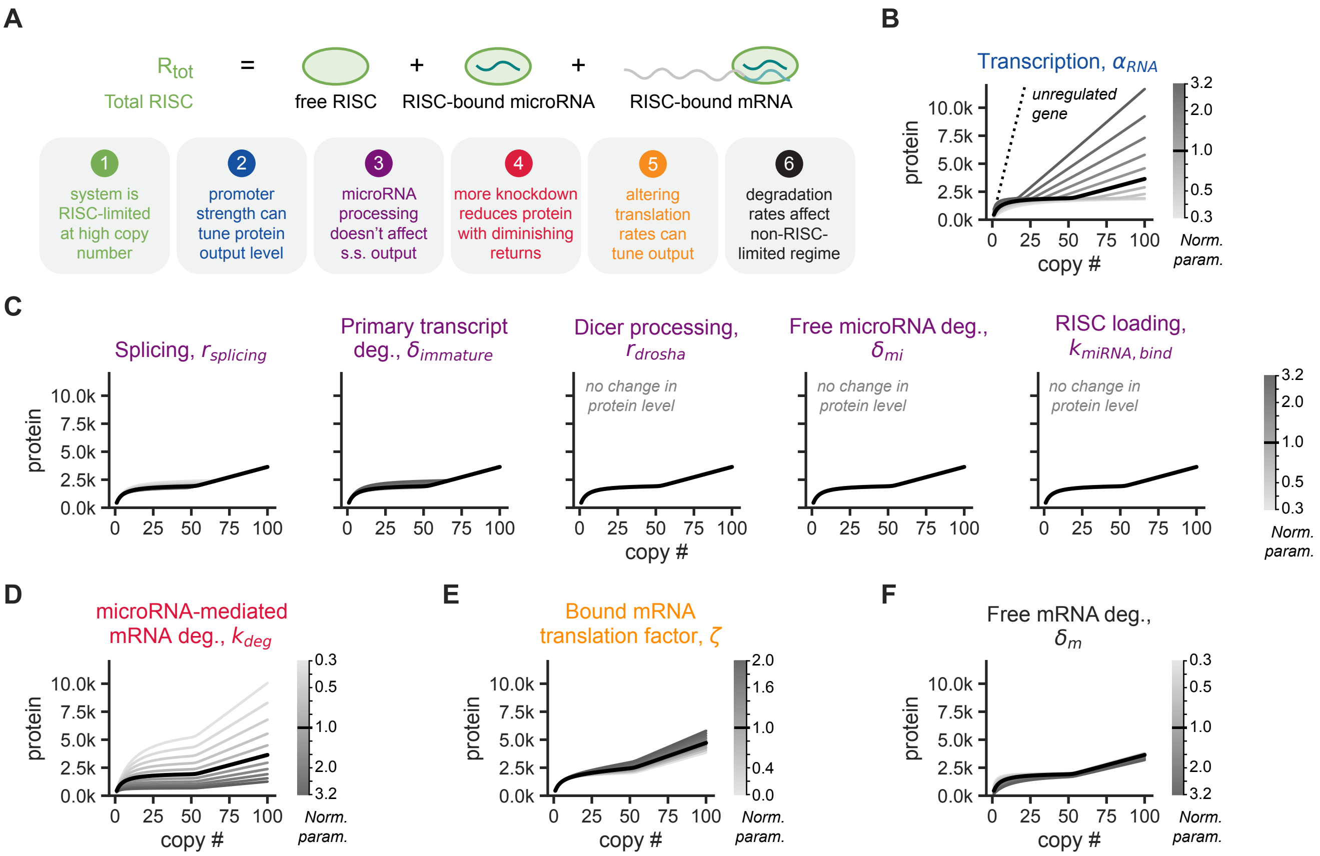


**Figure S2. Additional model parameter sweeps.**
**A**. Top: The total amount of RISC in the system, $R_{\mathrm{tot}}$, is the sum of the amount of free RISC, RISC bound to microRNA, and RISC-microRNA bound to mRNA. The total amount of RISC remains constant. Bottom: The six design principles revealed by the model, as illustrated in Figure 3.
**B**. Output protein level (in thousands of molecules) as a function of DNA copy number, $c_{\mathrm{regulated}}$, with a sweep of values of $\alpha_{\mathrm{RNA}}$, the transcription rate of the primary transcript. Chosen parameter values (shaded grey bar) are evenly log-distributed over an order of magnitude centered on the original parameter value. The black line in this panel and subsequent panels represents the output of the model with the original parameter values. Dashed black line indicates the solution for an unregulated gene.
**C**. Output protein level (in thousands of molecules) as a function of DNA copy number with a sweep of the values of the following parameters: splicing rate, $r_{\mathrm{splicing}}$; primary transcript degradation rate, $\delta_{\mathrm{immature}}$; Dicer processing rate, $r_{\mathrm{dicer}}$; free microRNA degradation rate, $\delta_{\mathrm{mi}}$; microRNA loading in RISC, $k_{\mathrm{miRNA,bind}}$. Chosen parameter values (shaded grey bar) are evenly log-distributed over an order of magnitude centered on the original parameter value. Output curves that do not change as the give parameter changes are indicated on the plot.
**D**. Output protein level (in thousands of molecules) as a function of DNA copy number with a sweep of values of $k_{\mathrm{deg}}$, the microRNA-mediated mRNA degradation rate. Chosen parameter values (shaded grey bar) are evenly log-distributed over an order of magnitude centered on the original parameter value.
**E**. Output protein level (in thousands of molecules) as a function of DNA copy number with a sweep of $\zeta$, the RISC-bound mRNA translation factor. Chosen parameter values (shaded grey bar) are linearly spaced between zero and one, where zero represents no translation of the bound mRNA and one represents translation at a rate equivalent to that of free mRNA.
**F**. Output protein level (in thousands of molecules) as a function of DNA copy number with a sweep of values of $\delta_{\mathrm{m}}$, the free mRNA degradation rate. Chosen parameter values (shaded grey bar) are evenly log-distributed over an order of magnitude centered on the original parameter value.
See model schematic in Figure 3A. Exact reactions, steady state analysis, and description of parameter sweeps can be found in Supplemental Information Section 1.

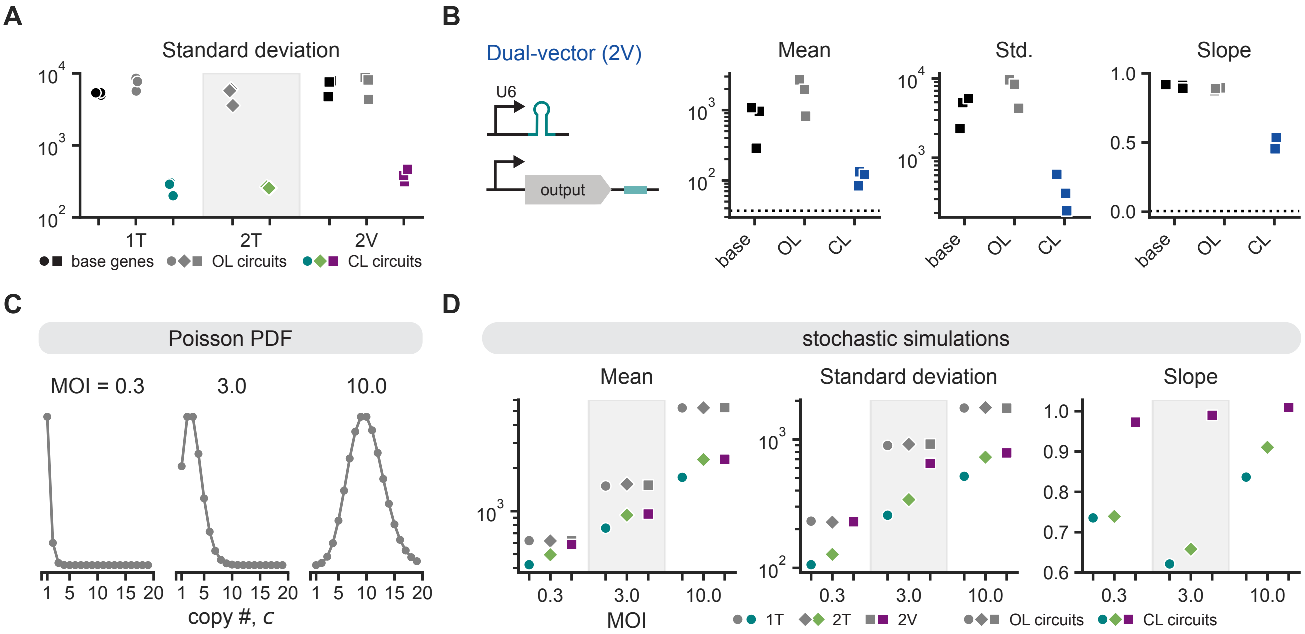


**Figure S3. Additional two-gene circuit implementations and simulations.**
**A**. Standard deviation of output expression for HEK293T cells transfected with base genes, closed-loop circuits, or open-loop circuits depicted in Figure 4A. All circuits use miRE-FF4 and are expressed by the EF1α promoter. Points represent individual biological replicates.
**B**. Left: DNA construct diagram of a two-vector circuit implementation with U6-driven microRNA expression. Right: Summary statistics of output expression for HEK293T cells transfected with the base gene, closed-loop circuit, or open-loop circuit depicted to the left. Presented mean values use the geometric mean. Std. refers to the standard deviation, and slope represents the slope of the line fitted to the binned, log-transformed marker-output points. Points represent individual biological replicates. Dashed lines represent values for cells transfected only with a marker gene lacking an intronic microRNA. All units are arbitrary units (AU) from a flow cytometer.
**C**. Probability density function (PDF) for Poisson distributions with means 0.3, 3, and 10 (left to right). These distributions are truncated such that all values are greater than or equal to one (i.e., the zero point of the PDF is removed). The distributions correspond to the MOI, or the effective "multiplicity of infection," in Figures S3D and 4D. The MOI is the mean of the Poisson distribution from which DNA copy number values, $c_{\text{regulated}}$ and $c_{\text{unregulated}}$, were chosen.
**D**. Summary statistics of 10,000 stochastic simulations run for each condition as described in Supplemental Information Section 1.2. Points represent protein geometric means (in molecules), standard deviations, and slopes for the distributions presented in Figure 4D.
1T, single-transcript; 2T, dual-transcript; 2V, dual-vector; base, base gene; OL, open-loop circuit; CL, closed-loop circuit.

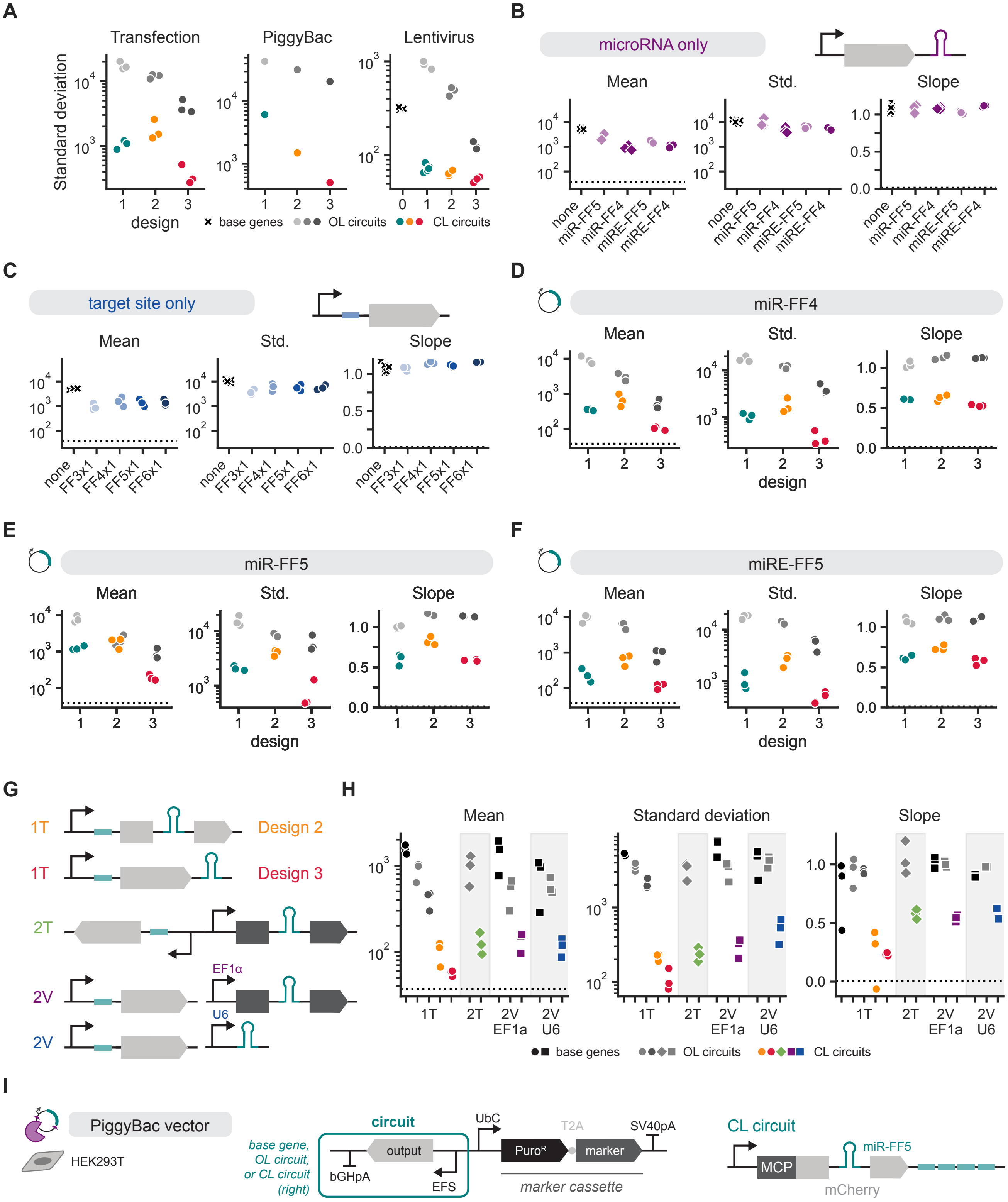
A
Transfection
PiggyBac
Lentivirus
Standard deviation
design
base genes
OL circuits
CL circuits
B
microRNA only
Mean
Std.
Slope
none
miR-FF5
miR-FF4
miRE-FF5
miRE-FF4
C
target site only
none
FF3x1
FF4x1
FF5x1
FF6x1
D
miR-FF4
E
miR-FF5
F
miRE-FF5
G
1T
Design 2
Design 3
2T
2V
EF1α
U6
H
Mean
Standard deviation
Slope
2V EF1a
2V U6
I
PiggyBac vector
HEK293T
base gene, OL circuit, or CL circuit (right)
circuit
output
bGHpA
EFS
UbC
T2A
SV40pA
Puro$^R$
marker
marker cassette
CL circuit
MCP
mCherry
miR-FF5

**Figure S4. Additional single-transcript and dual-transcript designs in transfection of HEK293T cells.**
**A**. Standard deviation of output expression in HEK293T cells transfected, PiggyBac-integrated, or lentivirally transduced with circuit designs as shown in Figure 5. Design "B" refers to the base gene.
**B**, **C**. Summary statistics for HEK293T cells transfected with output genes containing only a microRNA in the 3'UTR (S4B) or only a target site in the 5'UTR (S4C).
**D-F**. Summary statistics of output expression in HEK293T cells transfected with circuit designs in Figure 5 using miR-FF4 (S4D), miR-FF5 (S4E), or miRE-FF5 (S4F).
**G**. DNA construct diagrams for single-transcript (1T), dual-transcript (2T), and dual-vector (2V) circuits with microRNA target sites in the 5' UTR of the output gene.
**H**. Summary statistics of output expression in HEK293T cells transfected with the single-transcript, dual-transcript, or dual-vector circuits shown in Figure S4G.
**I**. DNA construct diagrams of PiggyBac vectors used for transposase-mediated integration in HEK293T cells. The output gene consists of an MCP-mCherry fusion protein, with miR-FF5 within an intron of the mCherry sequence. The closed-loop (CL) circuit contains four copies of matched (FF5) target sites, and the open-loop (OL) circuit contains four copies of orthogonal (FF3) target sites.
All data depicts expression of output genes driven by an EF1α promoter. The plotted mean values use the geometric mean. Std. refers to the standard deviation, and the slope represents the slope of the line fitted to the binned, log-transformed marker-output points. Dashed lines represent values for cells transfected only with a marker gene lacking an intronic microRNA. Points represent individual biological replicates. All units are arbitrary units (AU) from a flow cytometer.

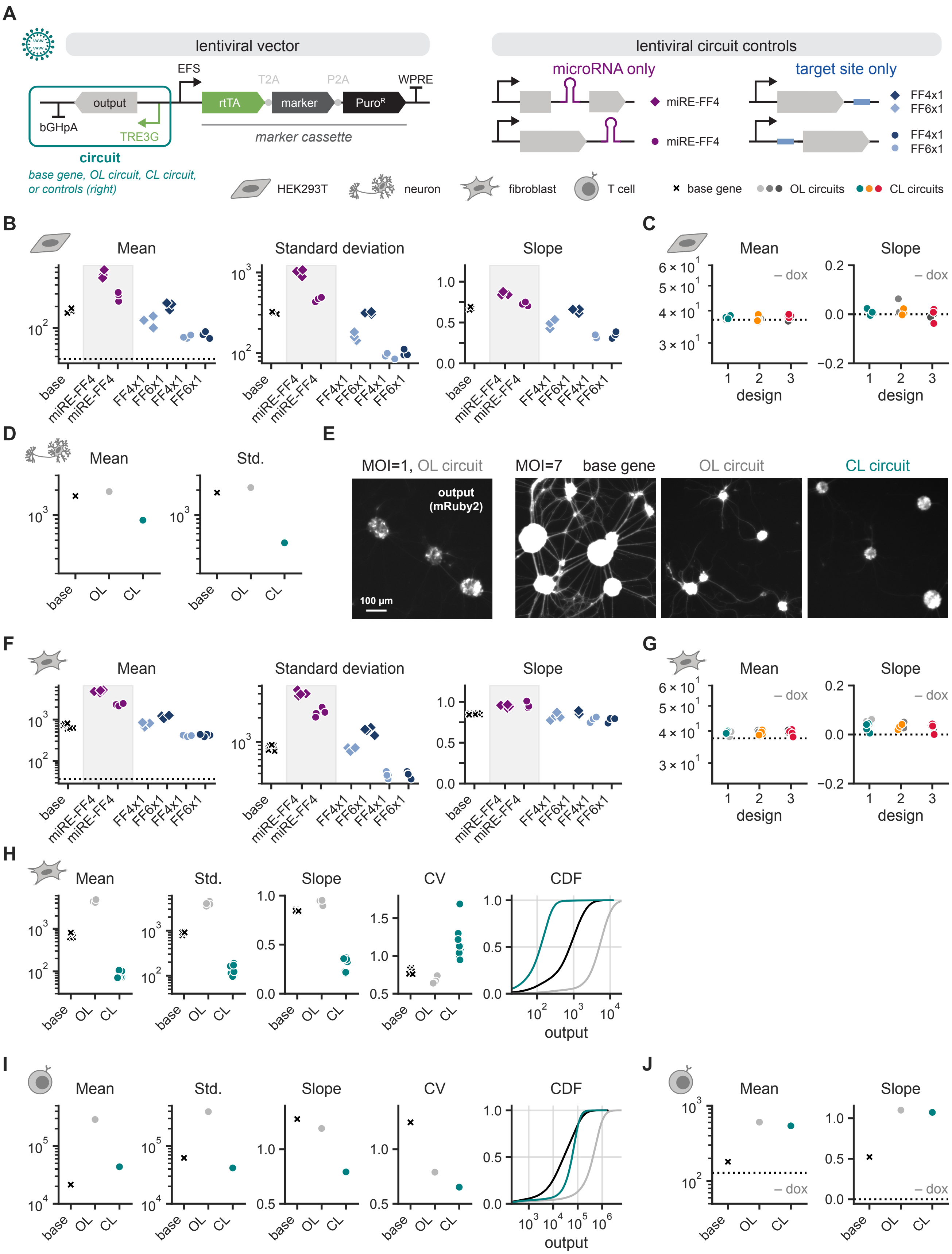
A
lentiviral vector
lentiviral circuit controls
output
bGHpA
TRE3G
EFS
rtTA
T2A
marker
P2A
PuroR
WPRE
marker cassette
circuit
base gene, OL circuit, CL circuit, or controls (right)
microRNA only
miRE-FF4
miRE-FF4
target site only
FF4x1
FF6x1
FF4x1
FF6x1
HEK293T
neuron
fibroblast
T cell
base gene
OL circuits
CL circuits
B
Mean
Standard deviation
Slope
base
miRE-FF4
miRE-FF4
FF4x1
FF6x1
FF4x1
FF6x1
C
Mean
Slope
– dox
design
D
Mean
Std.
base
OL
CL
E
MOI=1, OL circuit
MOI=7
base gene
OL circuit
CL circuit
output (mRuby2)
100 µm
F
Mean
Standard deviation
Slope
G
Mean
Slope
– dox
design
H
Mean
Std.
Slope
CV
CDF
output
I
Mean
Std.
Slope
CV
CDF
output
J
Mean
Slope
– dox

**Figure S5. Additional data and controls for lentiviral delivery to primary cells.**
**A**. Left: DNA construct diagram of the lentiviral vector used in transduction of primary cells. The output gene is placed on the antisense strand of the viral genome under the control of a doxycycline-inducible TRE3G promoter. The marker cassette—consisting of rtTA, a marker gene, and a puromycin-resistance gene separated by 2A peptides—is expressed divergently from an EFS promoter. Right: DNA construct diagrams for unregulated circuit controls consisting of the output gene with only either a microRNA or target site.
**B**. Output expression for HEK293T cells lentivirally transduced with unregulated control circuits in Figure S5A. Dashed line represents the mean of untransduced cells.
**C**. Output expression for HEK293T cells lentivirally transduced with open-loop or closed-loop circuits as in Figure 5D. Cells were cultured in the absence of the inducer doxycycline. Left dashed line represents the mean of untransduced cells, and right dashed line indicates a slope of zero.
**D**. Summary statistics of output expression for lentiviral transduction of primary rat cortical neurons as shown in Figure 6B.
**E**. Representative images of primary rat cortical neurons transduced with lentiviral vectors as in Figure 6B. Far left: Open-loop circuit transduced at an MOI of 1. Left, right, far right: Base gene, open-loop circuit, and closed-loop circuit, respectively, transduced at an MOI of 7. Images depict fluorescence of the output gene, mRuby2.
**F**. Output expression for primary mouse embryonic fibroblasts lentivirally transduced with unregulated control circuits in Figure S5A.
**G**. Output expression for primary mouse embryonic fibroblasts lentivirally transduced with open-loop or closed-loop circuits as in Figure 6E. Cells were cultured in the absence of the inducer doxycycline. Left dashed line represents the mean of untransduced cells, and right dashed line indicates a slope of zero.
**H**. Summary statistics of output expression for primary mouse embryonic fibroblasts lentivirally transduced with a base gene, open-loop circuit, or closed-loop circuit in the presence of doxycycline as in Figure 6E.
**I**. Summary statistics of output expression for primary human T cells lentivirally transduced with a base gene, open-loop circuit, or closed-loop circuit in the presence of doxycycline as in Figure 6F.
**J**. Output expression for primary human T cells lentivirally transduced with base gene, open-loop circuit, or closed-loop circuit as in Figure 6F. Cells were cultured in the absence of the inducer doxycycline. Left dashed line represents the mean of untransduced cells, and right dashed line indicates a slope of zero.
Base, base gene; OL, open-loop circuit; CL, closed-loop circuit. The plotted mean values use the geometric mean. Std. refers to the standard deviation, and the slope represents the slope of the line fitted to the binned, log-transformed marker-output points. CV is the coefficient of variation. CDF depicts the cumulative distribution function of output protein levels for one representative biological replicate. Points represent individual biological replicates. All units are arbitrary units (AU) from a flow cytometer.

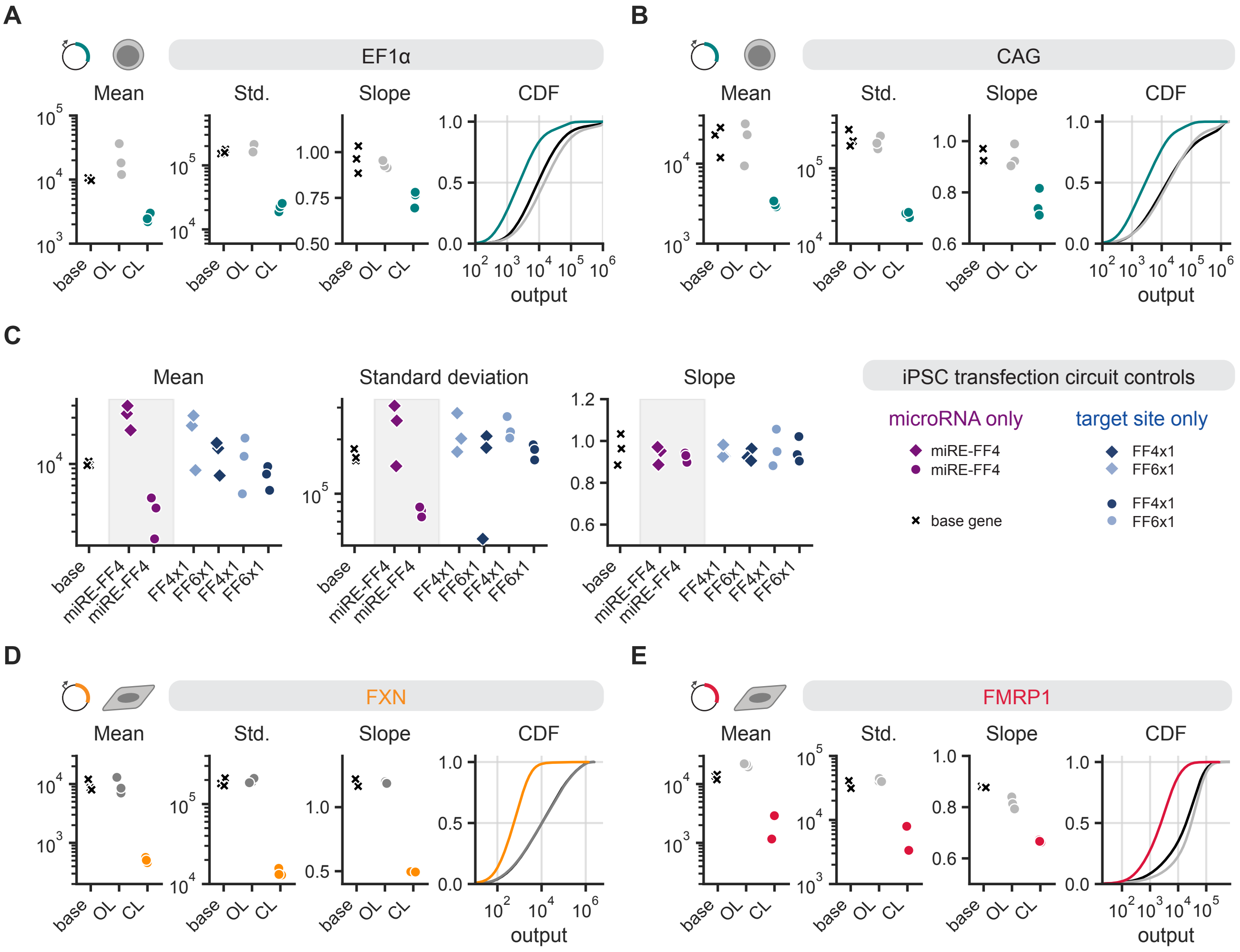


**Figure S6. Additional data and controls for iPSC transfection and transfection of therapeutically relevant genes.**
**A**, **B**. Summary statistics of output expression for induced pluripotent stem cells (iPSCs) co-transfected with a marker gene and a base gene, open-loop circuit, or closed-loop circuit expressed by the EF1α promoter (S6A, as in Figure 6G) or CAG promoter (S6B). Circuits use miRE-FF4.
**C**. Summary statistics of output expression for iPSCs transfected with unregulated circuit controls consisting of the output gene with only either a microRNA or target site.
**D**, **E**. Summary statistics of output expression for HEK293T cells transfected with circuits shown in Figure 6H regulating the therapeutically relevant genes FXN (Figure S6D) and FMRP1 (Figure S6E) as in Figure 6I.
Base, base gene; OL, open-loop circuit; CL, closed-loop circuit. The plotted mean values use the geometric mean. Std. refers to the standard deviation, and the slope represents the slope of the line fitted to the binned, log-transformed marker-output points. CDF depicts the cumulative distribution function of output protein levels for one representative biological replicate. Points represent individual biological replicates. All units are arbitrary units (AU) from a flow cytometer.

#### Supplementary Tables

| Figure | Channel | Laser (nm) | Filter | PMT Voltage |
|---|---|---|---|---|
| Figures 1, S1F to S1J, 2, 3, S3A, S3B, 4, S4A to S4F, S4H, 5B, 5D, 5E and 6E | mGL | 488 | 530 / 30 | 220 |
| | mRuby2 | 561 | 585 / 16 | 200 |
| Figure 6B | mGL | 488 | 530 / 30 | 220 |
| | mRuby2 | 561 | 585 / 16 | 260 |
| Figure 6I | EGFP | 488 | 530 / 30 | 220 |
| | mRuby2 | 561 | 615 / 25 | 300 |
| | iRFP | 637 | 670 / 14 | 320 |
| Figures S1B to S1E | mGL | 488 | 530 / 30 | 240 |
| | mRuby2 | 561 | 620 / 15 | 300 |
| Figures S4A and 5C | mGL | 488 | 530 / 30 | 220 |
| | mRuby2 | 561 | 585 / 16 | 360 |

**Table S1.** Attune channel assignments and settings are listed for all flow cytometry experiments except the T cell experiments.

| Figure | Channel | Laser (nm) | Filter | PMT Voltage |
|---|---|---|---|---|
| Figure 6F | mGL | 488 | 525 / 40 | 211 |
| | mRuby2 | 561 | 585 / 42 | 263 |
| | live-dead | 638 | 780 / 60 | 150 |

**Table S2.** Cytoflex S channel assignments and settings are listed for the flow cytometry T cell experiments.

### 1 Modeling

#### 1.1 Steady state analysis

We modeled the ComMAND system using the set of reactions shown in Figure 3A. Reactions were modeled as mass-action reactions with the stoichiometries and rate constants listed in Table S3.

| Reaction | Rate constant | $\text{mRNA}_i$ | pri | pre | miR | mRNA | R | RI | RIM | protein |
|---|---|---|---|---|---|---|---|---|---|---|
| Transcription | $c_{\text{regulated}}\alpha_{\text{RNA}}$ | 1 | 0 | 0 | 0 | 0 | 0 | 0 | 0 | 0 |
| Splicing | $r_{\text{splicing}}$ | -1 | 1 | 0 | 0 | 1 | 0 | 0 | 0 | 0 |
| Immature degradation | $\delta_{\text{immature}}$ | -1 | 0 | 0 | 0 | 0 | 0 | 0 | 0 | 0 |
| Drosha | $r_{\text{drosha}}$ | 0 | -1 | 1 | 0 | 0 | 0 | 0 | 0 | 0 |
| Dicer | $r_{\text{dicer}}$ | 0 | 0 | -1 | 1 | 0 | 0 | 0 | 0 | 0 |
| miRNA degradation | $\delta_{\text{miRNA}}$ | 0 | 0 | 0 | -1 | 0 | 0 | 0 | 0 | 0 |
| mRNA degradation | $\delta_{\text{mRNA}}$ | 0 | 0 | 0 | 0 | -1 | 0 | 0 | 0 | 0 |
| RISC loading | $k_{\text{miRNA,bind}}$ | 0 | 0 | 0 | -1 | 0 | -1 | 1 | 0 | 0 |
| Loaded degradation | $k_{\text{miRNA,deg}}$ | 0 | 0 | 0 | 0 | 0 | 1 | -1 | 0 | 0 |
| RISC-mRNA binding | $k_{\text{mRNA,bind}}$ | 0 | 0 | 0 | 0 | -1 | 0 | -1 | 1 | 0 |
| RISC-mRNA unbinding | $k_{\text{mRNA,unbind}}$ | 0 | 0 | 0 | 0 | 1 | 0 | 1 | -1 | 0 |
| mRNA knockdown | $k_{\text{deg}}$ | 0 | 0 | 0 | 0 | 0 | 0 | 1 | -1 | 0 |
| Translation | $\alpha_p[\text{mRNA}]$ | 0 | 0 | 0 | 0 | 0 | 0 | 0 | 0 | 1 |
| Bound translation | $\zeta\alpha_p[\text{RIM}]$ | 0 | 0 | 0 | 0 | 0 | 0 | 0 | 0 | 1 |
| Protein degradation | $\delta_p$ | 0 | 0 | 0 | 0 | 0 | 0 | 0 | 0 | -1 |

**Table S3.** The stoichiometry of our model reactions of the ComMAND architecture modeled, as shown in Figure 3A, is given, along with the rate constant. "miR" refers to the microRNA, "R" refers to unbound RISC, "RI" refers to RISC loaded with miRNA, and "RIM" refers to loaded RISC bound to mRNA. The rate of a reaction with reaction rate constant $r$ and stoichiometric coefficients $\zeta_i$ for species $s_i$ is $r\prod_{\zeta_i<0} s_i^{|\zeta_i|}$, e.g. a mass action equation in terms of the reactants (species with negative stoichiometric coefficients)

To analyze the behavior of the dual-transcript, dual-vector, and U6 systems presented in Figures 4 and S4G, we replaced the first two reactions ("Transcription", "Splicing") with the reactions listed in Table S4. To understand the system, we computed an analytical steady-state solution for this system.

| Architecture | Reaction | Rate constant | $\text{mRNA}_i$ | pri | mRNA | $\text{mRNA}_{\text{ignore}}$ |
|---|---|---|---|---|---|---|
| Dual-transcript (Figure 4) | miRNA transcription | $c_{\text{regulated}}\alpha_{\text{RNA}}$ | 1 | 0 | 0 | 0 |
| | Splicing | $r_{\text{splicing}}$ | -1 | 1 | 0 | 1 |
| | Regulated transcription | $c_{\text{regulated}}\alpha_{\text{RNA}}$ | 0 | 0 | 1 | 0 |
| Dual-vector (Figure 4) | miRNA transcription | $c_{\text{regulated}}\alpha_{\text{RNA}}$ | 1 | 0 | 0 | 0 |
| | Splicing | $r_{\text{splicing}}$ | -1 | 1 | 0 | 1 |
| | Regulated transcription | $c_{\text{unregulated}}\alpha_{\text{RNA}}$ | 0 | 0 | 1 | 0 |
| U6 (Figure S4G) | miRNA transcription | $c_{\text{U6}}\alpha_{\text{U6}}$ | 0 | 1 | 0 | 0 |
| | Regulated transcription | $c_{\text{regulated}}\alpha_{\text{RNA}}$ | 0 | 0 | 1 | 0 |

**Table S4.** Alternate reactions used to simulate the dual-transcript, dual-vector, and dual-vector U6 architectures are given as a stoichiometry table. In the dual-transcript and dual-vector cases, splicing creates an unregulated output transcript, $\text{mRNA}_{\text{ignore}}$ that does interact with the rest of the system. The dual-transcript and dual-vector cases then only differ in the copy number variable used in the transcription reactions. For the dual-vector U6 architecture, the transcription steps directly create pri-microRNA or the regulated transcript.

##### 1.1.1 ComMAND (Single-transcript)

For the single-transcript case, we have the following ODEs defining the production of the pre-mRNA:

$$\begin{aligned}
\frac{\mathrm{d}\text{mRNA}_i(t)}{\mathrm{d}t} &= -\, r_{\text{splicing}}\text{mRNA}_i(t) + c_{\text{regulated}}\alpha_{\text{RNA}} - \text{mRNA}_i(t)\delta_{im} \\
\frac{\mathrm{d}\text{pri}(t)}{\mathrm{d}t} &= -\, r_{\text{drosha}}\text{pri}(t) + r_{\text{splicing}}\text{mRNA}_i(t) \\
\frac{\mathrm{d}\text{pre}(t)}{\mathrm{d}t} &= -\, r_{\text{dicer}}\text{pre}(t) + r_{\text{drosha}}\text{pri}(t)
\end{aligned}$$

At steady state, this can be solved:

$$\text{mRNA}_i = \frac{c_{\text{regulated}}\alpha_{\text{RNA}}}{\delta_{im} + r_{\text{splicing}}} \quad (1)$$

$$\text{pri} = \frac{r_{\text{splicing}}}{r_{\text{drosha}}} \cdot \frac{c_{\text{regulated}}\alpha_{\text{RNA}}}{\delta_{im} + r_{\text{splicing}}} \quad (2)$$

$$\text{pre} = \frac{r_{\text{splicing}}}{r_{\text{dicer}}} \cdot \frac{c_{\text{regulated}}\alpha_{\text{RNA}}}{\delta_{im} + r_{\text{splicing}}} \quad (3)$$

The effective production rate of the controlled mRNA is, at steady state:

$$\tilde{r}_{\text{controlled}} = r_{\text{splicing}}\text{mRNA}_i \quad (4)$$

##### 1.1.2 Dual-transcript

In the dual transcript case, the regulated mRNA is unspliced, so the only immature mRNA species is the one containing the miRNA. From this intronic transcript, the pri-microRNAs are created.

The equations to solve at steady state are:

$$\frac{\mathrm{dmRNA}_i(t)}{\mathrm{d}t} = - r_{\text{splicing}}\mathrm{mRNA}_i(t) + c_{\text{unregulated}}\alpha_{\text{RNA}} - \mathrm{mRNA}_i(t)\delta_{im}$$
$$\frac{\mathrm{dpri}(t)}{\mathrm{d}t} = - r_{\text{drosha}}\mathrm{pri}(t) + r_{\text{splicing}}\mathrm{mRNA}_i(t)$$
$$\frac{\mathrm{dpre}(t)}{\mathrm{d}t} = - r_{\text{dicer}}\mathrm{pre}(t) + r_{\text{drosha}}\mathrm{pri}(t)$$

which at steady state give

$$\mathrm{pri} = \frac{r_{\text{splicing}}}{r_{\text{drosha}}} \cdot \frac{c_{\text{unregulated}}\alpha_{\text{RNA}}}{\delta_{im} + r_{\text{splicing}}} \tag{5}$$
$$\mathrm{pre} = \frac{r_{\text{splicing}}}{r_{\text{dicer}}} \cdot \frac{c_{\text{unregulated}}\alpha_{\text{RNA}}}{\delta_{im} + r_{\text{splicing}}} \tag{6}$$

Here, the effective production rate is directly dependent on the copy number and the transcription rate:

$$\tilde{r}_{\text{controlled}} = c_{\text{regulated}}\alpha_{\text{RNA}} \tag{7}$$

###### 1.1.3 Dual-vector

In the dual-vector case, the steady-state solution is the same as in Equations (5) and (6). However, the effective production rate of the controlled transcript depends on a different copy number

$$\tilde{r}_{\text{controlled}} = c_{\text{unregulated}}\alpha_{\text{RNA}} \tag{8}$$

###### 1.1.4 Non-intronic dual transcript

In this simplest case, there is no splicing involved with the production of the miRNA. Instead, we have the equations:

$$\frac{\mathrm{dpri}(t)}{\mathrm{d}t} = - r_{\text{drosha}}\mathrm{pri}(t) + c_{\text{U6}}\alpha_{\text{U6}}$$
$$\frac{\mathrm{dpre}(t)}{\mathrm{d}t} = - r_{\text{dicer}}\mathrm{pre}(t) + r_{\text{drosha}}\mathrm{pri}(t)$$

which can be solved at steady state

$$\mathrm{pri} = \frac{c_{\text{U6}}\alpha_{\text{U6}}}{r_{\text{drosha}}} \tag{9}$$
$$\mathrm{pre} = \frac{c_{\text{U6}}\alpha_{\text{U6}}}{r_{\text{dicer}}} \tag{10}$$

The effective production rate of controlled mRNA is:

$$\tilde{r}_{\text{controlled}} = c_{\text{regulated}}\alpha_{\text{RNA}} \tag{11}$$

##### 1.1.5 Shared system

All implementations of the miR-iFFL share a common set of reactions, as listed below the line in Table S3. We can think of the set of common reactions as a network that takes in two species—the *controlled/regulated mRNA* and the *pre-miRNA*—and outputs a final protein (and mRNA) concentration.

The specific form of the regulated mRNA term differs between each architecture, so we use the effective production rate $\tilde{r}_{\text{controlled}}$ computed earlier.

These reactions are:

$$\frac{\text{miR}(t)}{\text{d}t} = r_{\text{dicer}}\text{pre}(t) - \text{miR}\,(t)\,\delta_{mi} - k_{miR\_b}\text{miR}\,(t)\,\text{R}(t) \tag{12}$$

$$\frac{\text{dR}(t)}{\text{d}t} = k_{miR\_deg}\text{RI}(t) - k_{miR\_b}\text{miR}\,(t)\,\text{R}(t) \tag{13}$$

$$\begin{aligned}\frac{\text{dRI}(t)}{\text{d}t} =& + k_{deg}\text{RIM}(t) + k_{mRNA\_ub}\text{RIM}(t) - k_{mRNA\_b}\text{RI}(t)\text{mRNA}(t) \\ & - k_{miR\_deg}\text{RI}(t) + k_{miR\_b}\text{miR}\,(t)\,\text{R}(t)\end{aligned} \tag{14}$$

$$\frac{\text{dRIM}(t)}{\text{d}t} = - k_{deg}\text{RIM}(t) - k_{mRNA\_ub}\text{RIM}(t) + k_{mRNA\_b}\text{RI}(t)\text{mRNA}(t) \tag{15}$$

$$\frac{\text{dmRNA}(t)}{\text{d}t} = k_{mRNA\_ub}\text{RIM}(t) + \tilde{r}_{\text{controlled}} - \delta_m\text{mRNA}(t) - k_{mRNA\_b}\text{RI}(t)\text{mRNA}(t) \tag{16}$$

$$\frac{\text{d}P(t)}{\text{d}t} = - \delta_p P(t) + \alpha_p\text{mRNA}(t) + \zeta\alpha_p\text{RIM}(t) \tag{17}$$

Immediately, we can solve Equation (12) and Equation (17) at steady state to get:

$$\text{miR} = \frac{r_{\text{dicer}} \cdot \text{pre}}{k_{miR\_b} \cdot \text{R} + \delta_{mi}} \tag{18}$$

$$P = \frac{\alpha_p}{\delta_p}(\text{mRNA} + \zeta \cdot \text{RIM}) \tag{19}$$

To solve the rest at steady state, note that equations Equations (13) to (15) sum to zero at steady state; this means that we will need to introduce the RISC conservation equation and substitute to fully solve this system:

$$R_{\text{tot}} = \text{R} + \text{RI} + \text{RIM} \tag{20}$$

We can then solve for RI using Equation (13):

$$\text{RI} = \frac{k_{miR\_b}r_{\text{dicer}} \cdot \text{pre} \cdot \text{R}}{k_{miR\_ub}(\delta_{mi} + k_{mi\_b} \cdot \text{R})} = \frac{R}{\kappa_1 + \kappa_2 R} \tag{21}$$

for

$$\kappa_1 = \frac{k_{miR\_ub}\delta_{mi}}{k_{miR\_b}r_{\text{dicer}} \cdot \text{pre}} \qquad \kappa_2 = \frac{k_{miR\_ub}k_{mi\_b}}{k_{miR\_b}r_{\text{dicer}} \cdot \text{pre}}$$

Similarly, by plugging this result back in and using Equation (15), we can define:

$$\kappa_3 = \frac{\delta_m\delta_{mi}k_{miR\_ub}(k_{\text{deg}} + k_{mRNA\_ub})}{k_{mRNA\_b}k_{miR\_b}r_{\text{dicer}} \cdot \text{pre} \cdot \tilde{r}_{\text{controlled}}}$$

$$\kappa_4 = \frac{\delta_m k_{miR\_ub}(k_{\text{deg}} + k_{mRNA\_ub})}{k_{mRNA\_b} r_{\text{dicer}} \cdot \text{pre} \cdot \tilde{r}_{\text{controlled}}} + \frac{k_{\text{deg}}}{\tilde{r}_{\text{controlled}}}$$

so that

$$\text{RIM} = \frac{R}{\kappa_3 + \kappa_4 R} \tag{22}$$

Then, the RISC mass closure equation can be written:

$$\begin{aligned} R_{\text{tot}} &= \text{R} + \frac{\text{R}}{\kappa_1 + \kappa_2 \text{R}} + \frac{\text{R}}{\kappa_3 + \kappa_4 \text{R}} \\ 0 &= (\text{R} - R_{\text{tot}})(\kappa_1 + \kappa_2 \text{R})(\kappa_3 + \kappa_4 \text{R}) + (\kappa_3 + \kappa_4 \text{R})\text{R} + (\kappa_1 + \kappa_2 \text{R})\text{R} \end{aligned}$$

This is a cubic equation in R:

$$0 = (\kappa_2\kappa_4)\text{R}^3 + (\kappa_1\kappa_4 + \kappa_2\kappa_3 + \kappa_4 + \kappa_2 - R_{\text{tot}}\kappa_2\kappa_4)\text{R}^2 + (\kappa_1\kappa_3 + \kappa_3 + \kappa_1 - R_{\text{tot}}(\kappa_1\kappa_4 + \kappa_2\kappa_3))\text{R} - R_{\text{tot}}\kappa_1\kappa_3 \tag{23}$$

While this represents an analytic steady state solution for R, it is only tractable when solved numerically. For the parameter regimes chosen here, the cubic discriminant is positive, which means that there are three real roots. In practice—and at every parameter range included in this work—two of the roots are unphysical (negative), so we choose the positive root.

With this positive root, we can solve for the steady state mRNA concentration:

$$\text{mRNA} = \tilde{r}_{\text{controlled}} \left( \delta_m + \frac{k_{\text{deg}} k_{mRNA\_b} k_{miR\_b} r_{\text{dicer}} \cdot \text{pre} \cdot \text{R}}{k_{miR\_ub}(k_{\text{deg}} + k_{mRNA\_ub})(\delta_{mi} + k_{miR\_b} \cdot \text{R})} \right)^{-1} \tag{24}$$

By back-substituting this result into Equation (19) and Equation (22), we can compute the steady-state protein concentration.

#### 1.2 Stochastic Simulations

We first constructed models of the dual-transcript and dual-vector architectures, in which the output mRNA is directly transcribed rather than spliced from a primary transcript containing the microRNA (Figure 4C and Table S4). In the dual-transcript case, the DNA copy number of the output gene is the same as that of the microRNA gene, since they are physically located on the same plasmid. For the dual-vector circuit, the DNA copy number of the output gene is allowed to vary from the that of the microRNA gene, since co-delivery of two vectors may not be perfect. A steady-state analytical solution for these systems does not provide much insight into differences between the architectures, since it does not account for effects of noise on copy number or output production.

Therefore, we performed stochastic simulations of the reaction systems for each circuit architecture. We modeled the systems at three different effective multiplicity of infections (MOIs), a common parameter in viral transductions that specifies the number of viral particles added per cell. In this context, this copy number can be modeled as a Poisson distribution with a mean equal to the MOI. To recapitulate this variation in delivery in our model, we chose DNA copy numbers for each simulation run from a Poisson distribution with one of three different means: 0.3, to simulate single-copy integration; 3, to simulate a low-copy regime; and 10, to simulate a high-copy regime (Figure S3C).

The resulting systems were simulated using a Gillepse algorithm implementation (Cataylst.jl, DifferentialEquations.jl, and JumpProblems.jl).

#### 1.3 Parameter values

For the steady-state and stochastic simulations, we used the following parameter values:

| Parameter | Value | Units |
|---|---|---|
| $\alpha_{\text{RNA}}$ | $4.62 \cdot 10^{-2}$ | 1 / s |
| $r_{\text{splicing}}$ | $2.0 \cdot 10^{-3}$ | 1 / s |
| $\delta_{\text{immature}}$ | $2.88 \cdot 10^{-4}$ | 1 / s |
| $r_{\text{drosha}}$ | $1.0 \cdot 10^{-2}$ | 1 / s |
| $r_{\text{dicer}}$ | $1.0 \cdot 10^{-3}$ | 1 / s |
| $\delta_{\text{miRNA}}$ | $2.88 \cdot 10^{-4}$ | 1 / s |
| $\delta_{\text{mRNA}}$ | $2.88 \cdot 10^{-4}$ | 1 / s |
| $k_{\text{miRNA,bind}}$ | $1.0 \cdot 10^{-5}$ | 1 / (molecules · s) |
| $k_{\text{miRNA,deg}}$ | $2.16 \cdot 10^{-5}$ | 1 / s |
| $k_{\text{mRNA,bind}}$ | $1.84 \cdot 10^{-6}$ | 1 / (molecules · s) |
| $k_{\text{mRNA,unbind}}$ | 0.303 | 1 / s |
| $k_{\text{deg}}$ | $7.0 \cdot 10^{-3}$ | 1 / s |
| $\alpha_p$ | $3.33 \cdot 10^{-4}$ | 1 / s |
| $\zeta$ | 0.0 | dimensionless |
| $\delta_p$ | $9.67 \cdot 10^{-5}$ | 1 / s |

**Table S5.** Base parameter values used in the model. For Figure 3, parameter values are varied from these base values while keeping non-modified parameters constant.
